# Coexistence of lipid phases stabilizes interstitial water in the outer layer of mammalian skin

**DOI:** 10.1101/786699

**Authors:** Christopher M. MacDermaid, Kyle Wm. Hall, Russell H. DeVane, Michael L. Klein, Giacomo Fiorin

## Abstract

The lipid matrix in the outer layer of mammalian skin, the stratum corneum, has been previously investigated by multiple biophysical techniques, aimed at identifying hydrophilic and lipophilic pathways of permeation. While consensus is developing over the microscopic structure of the lipid matrix, no molecular-resolution model describes the permeability of all chemical species simultaneously. Using molecular dynamics simulations of a model mixture of skin lipids, the self-assembly of the lipid matrix lamellae has been studied. At higher humidity, the resulting lamellar phase is maintained by partitioning excess water into isolated droplets of controlled size and spatial distribution. The droplets may fuse together to form intra-lamellar water channels, thereby providing a pathway for the permeation of hydrophilic species. These results reconcile competing data on the outer skin’s structure and broaden the scope of molecular-based methods to improve the safety of topical products and to advance transdermal drug delivery.

## INTRODUCTION

The *stratum corneum* of mammalian skin (SC) provides the outermost and primary barrier against harmful chemicals and pathogens (Elias, 2005). Circumventing its function in a controlled manner has allowed for the transdermal delivery of drugs to treat a broad range of medical conditions (Hirvonen et al., 1996). There is a demonstrated need to further understand the skin’s permeability mechanisms, which can be measured quantitatively (Pham et al., 2017) but not spatially resolved at the molecular level. Cells in the SC, called *corneocytes*, are almost entirely made up by a solid keratin mesh with water amounting to less than 50% of their dry weight (Caspers et al., 2001). A *lipid matrix* about 200 nm-thick envelops the corneocytes and fills the interstices between them. The lipid matrix has been the object of nanometer-resolution studies by many laboratories (Bouwstra et al., 2001; Das et al., 2013; Del Regno & Notman, 2018; Gupta et al., 2016; Iwai et al., 2012; McIntosh et al., 1996; Moore et al., 1997; Moore et al., 2018; Schroter et al., 2009; Skolova et al., 2013; Wang & Klauda, in press; White et al., 1988).

Unlike the plasma membranes of viable cells, the SC’s lipid matrix must minimize permeability but also adapt to the mechanical strain and to the changes in humidity and pH associated with epidermal growth. This function is achieved by a mixture of lipid species, whose primary constituents are ceramides, cholesterol and free fatty acids in comparable proportions. Due to the absence of electron-dense phosphate groups and the very low humidity of the lipid matrix, biophysical detection methods are limited in resolution and significant uncertainty remains. The most widely accepted fact is that the lipid matrix is made up of multiple lamellae, lying parallel to the nearest corneocyte’s surface (Hou et al., 1991).

A single bilayer of skin lipids forms the so-called *short-periodicity phase* (SPP), approximately 5.4 nm in thickness. However, the typical thickness of the lipid lamellae in extracted skin samples is about 13 nm (Hou et al., 1991), consistent with the *long-periodicity phase* (LPP) observed in mixtures of skin lipids (Bouwstra et al., 1996). The LPP and the SPP are often detected in coexistence with each other, but the relative stability of the two is not established universally (Schmitt et al., 2019), nor is that of other reported thicknesses (Schroter et al., 2009).

To explain the LPP, a trilayer or “sandwich” model has been proposed (Beddoes et al., 2018; Bouwstra et al., 2001; Mojumdar et al., 2015a), wherein two liquid-ordered bilayers are thought to enclose a central liquid-disordered layer through which permeation may occur. NMR experiments also showed that the skin lipids are in equilibrium between “solid” and “liquid” states, with the “solid” state being dominant (Pham et al., 2017). Due to the long times (several hours) required for the LPP to emerge in synthetic samples, it is currently not possible to use computation to quantify the relative stability of each phase.

Aside from lamellae, bicontinuous cubic and other negative-curvature phases have also been reported by cryo-microscopy imaging of the SC at different stages of development (Norlen, 2001) and molecular dynamics (MD) simulations (Das et al., 2013). These results correlate well with the intrinsic preference of ceramides for negative curvature (Veiga et al., 1999) and it has been suggested that the skin lipids only form lamellar phases when confined between the rigid corneocytes (Das et al., 2013). According to this hypothesis, lamellar phases would be obtained *in vitro* only through the use of a heating process targeted at reproducing the *in vivo* structure (Bouwstra et al., 1996).

The most significant limitation of the models described above, particularly the lamellar phases, is that they do not explain all available experimental permeability data. For small lipophilic molecules, permeability measurements from skin samples led to the development of empirical models (Potts & Guy, 1992) whose accuracy is comparable to atomistic computations (Lundborg et al., 2018). On the other hand, hydrophilic molecules and macromolecules are found to permeate through the skin many orders of magnitude more quickly than through lipid bilayers (Mitragotri et al., 2011). It has been hypothesized by an increasing number of researchers that these molecules exploit an alternative *hydrophilic pathway*, formed by sporadic *water channels* of microscopic diameter running through the SC’s lipid matrix (Flynn, 1989; Menon & Elias, 1997; Mitragotri et al., 2011). Although evidence exists that permeation of water is inhomogeneous through the lipid matrix (Menon & Elias, 1997), the presence of nanometer-size channels and their mechanism of formation have not been established experimentally.

The simulation results presented in this paper show that the lipid phases described above can coexist at close distance. Bilayers of a skin lipid mixture rich in long-chain ceramides are shown to contain a central liquid-disordered region, a property that is shared with the LPP under the “sandwich” model. Permeation across such interface yields accurate predictions of the permeability of lipophilic molecules through the skin. Stable lamellae with LPP thickness are also formed via the spontaneous demixing of individual lipids, which follow with good approximation the distributions modeled from neutron diffraction (Beddoes et al., 2018; Mojumdar et al., 2015a). Such comparison is free from bias, because measurements of the LPP are not used in the simulation energy function and the initial configurations are randomized.

Also shown in this study is that when water is allowed to diffuse into the lamellae, it aggregates into inverse-micellar droplets surrounded by lipid head groups; despite significant humidity in the outer water buffer, these droplets account only for a 1.7 water:lipid ratio in the lamellar interior. The size and shape of the droplets are quantitatively described by a mechanical model based on experimental data. The same model predicts the formation of channels as a rare event, with activation energy well within the range that can be supplied by permeation enhancers. These results show that the ability of the skin’s lipids to appear both in “solid” and “liquid” states (Bouwstra et al., 2001; Pham et al., 2017) explains the simultaneous presence of multiple permeability pathways for distinct chemical species.

## RESULTS

The first three Results sections discuss simulations of a model mixture of skin lipids in the bilayer phase: *(i)* self-assembled models from coarse-grained (CG) simulations are compared to atomistic and experimental data; *(ii)* permeabilities of small molecules across the atomistic SPP bilayer model are also computed, and *(iii)* it is demonstrated that heating of bilayers induces the confinement of water as droplets or channels. In the following sections, self-assembled lamellae similar to the LPP are shown to contain water droplets that are formed by spontaneous aggregation of individual molecules at physiological temperature. As a control, the formation of dehydrated lamellae is also simulated, showing fair agreement with the LPP. Lastly, the size distribution of the droplets and its implications on the formation of continuous channels are discussed.

### Self-assembly of model skin lipid mixtures and computational model validation

Model mixtures of the primary components of the lipid matrix of the stratum corneum (ceramides, cholesterol and free fatty acids) were characterized by atomistic and coarse-grained (CG) molecular dynamics (MD) simulations. Two ceramide species were included, *CER[EOS]* with a C30:0 fatty acid chain that is ester-linked to a C18:2 linoleic acid chain, and *CER[NS]* with a C24:0 fatty acid chain (**Fig. 1**). Models that include free fatty acids (FFAs) were prepared using behenic acid (C22:0), which is at the peak of the FFA chain length distribution in the SC (Wertz et al., 1991).

**Figure 1.**
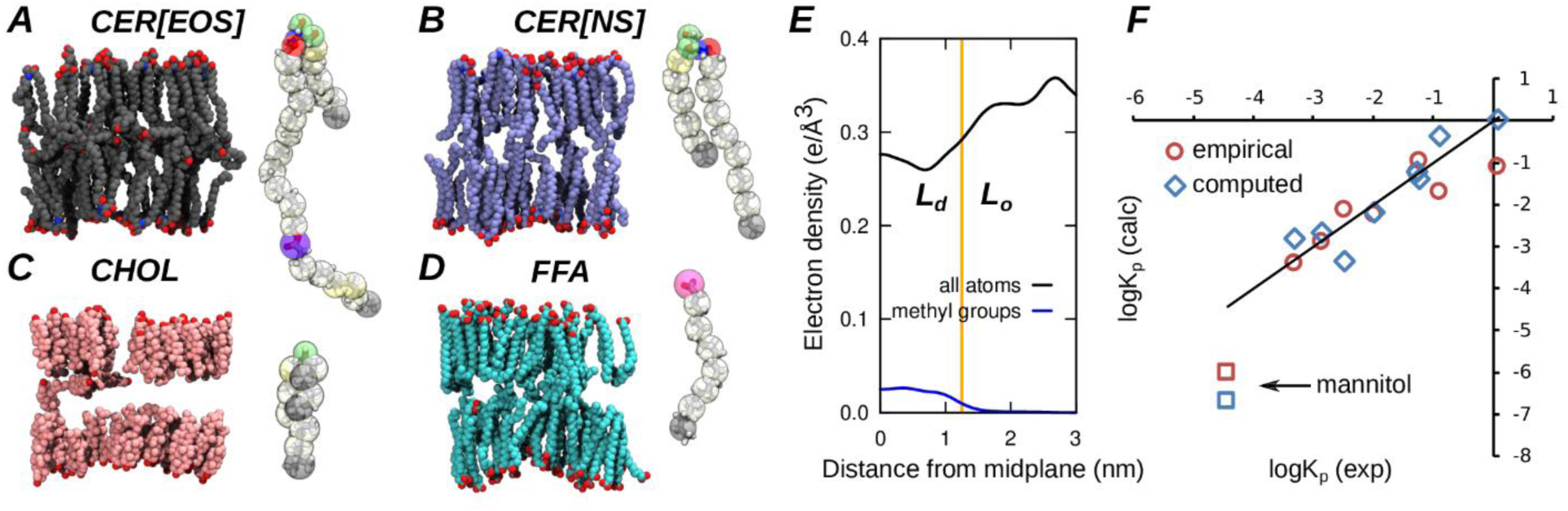
Structure and permeability of skin lipid model bilayers. Panels **A-D** show a bilayer snapshot for each of the lipids used in this study (atomistic detail, hydrogens hidden): CER[EOS] (A), CER[NS] (B), cholesterol (C), and behenic acid (D) are colored in gray, blue, pink and cyan respectively. To the right of each snapshot is a schematic of the coarse-grained (CG) representation of the molecule. Panel **E** shows the electron density profile (EDP) of the bilayer (black) and that of terminal methyl groups (blue); the orange line is the boundary between liquid- ordered (L_o_) and liquid-disordered (L_d_) regions. Panel **F** compares the computed skin permeability *k*_P_ (blue) with empirical estimates from the Potts-Guy equation (red); squares indicate values for mannitol. Root-mean-squared errors (RMSEs) on log(*k*_P_) are 0.73 and 0.72 for the computed and empirical values, respectively.

The CG simulation model was tested on simple mixtures of the two ceramides by characterizing the self-assembly of models of different sizes from randomized initial conditions (**Fig. S1**). A 5:1 water:lipid ratio was used, which is higher than the 1:1 or 2:1 water:lipid ratios inferred from neutron diffraction (Groen et al., 2011; Schroter et al., 2009). Such conditions ensured that results are transferable between atomistic simulations, which treat water molecules independently, and CG simulations, which treat three water molecules as a single CG particle that does not form ordered hydrogen networks.

Mixtures of ceramides, cholesterol and water self-assembled consistently into lamellar phases within hundreds of nanoseconds of CG-MD simulation (**Fig. S1**). Bilayers were also simulated in atomistic and CG resolution, and both compared to experimental data. Simulations of CER[NS] exhibit gel- or liquid-lamellar phases depending on the computational model used, and give rise to distinct spacings (**Fig. S2**) that correspond to lamellar diffraction peaks measured for a ceramide mixture with CER[NS] as the primary component (Mojumdar et al., 2015b).

In the presence of cholesterol, the atomistic and CG computational models of both CER[EOS] and CER[NS] deliver very similar bilayer thickness, hexatic order parameter and lipid distribution (**Figs. S2-S5**). Thicknesses are in agreement with diffraction measurements: about 7.5 nm for CER[EOS]:CHOL vs. 7.7 nm (Groen et al., 2010), and about 5.4 nm for CER[NS]:CHOL vs. 5.36 nm of a mixture with predominantly CER[NS] (Groen et al., 2011). In all simulations the ceramide head groups are in contact with the water layer; the sphingosine chain, the fatty acid chain of CER[NS] (C24:0), and the saturated portion (C30:0) of the CER[EOS] fatty acids chains are highly ordered (**Fig. S6**). Instead, the ester-linked unsaturated chains (C18:2) of CER[EOS] are in a liquid-disordered isotropic configuration with a significant fraction of “hooked” conformations, consistent with nuclear-magnetic-resonance measurements (Pham et al., 2018). The same results apply to the model mixture of CER[EOS], CER[NS], cholesterol and fatty acid in a 1:1:2:2 ratio (**Fig. 1**), used to model the lipid matrix.

In macroscopic samples of skin lipids, ordered lamellar phases like the SPP and the LPP are not easily formed via self-assembly at room temperature. However, as hypothesized earlier (Das et al., 2013) the assembly of relatively large model systems into lamellar structures can also be facilitated by the presence of a template (**Fig. S7**). Due to the use of water in the mixture and the intrinsic preference of the CG model for hydrated interfaces, only the bilayer phase (SPP) is observed. In samples of similar composition to the one here used, the SPP is competitive with the LPP. The SPP can be suppressed only by using higher concentrations of CER[EOS] (Beddoes et al., 2018) even when applying heat above 80 °C followed by several hours of equilibration at room temperature (a procedure used to enhance the rate of formation of the LPP). Except for the atomistic simulations shown in **Fig. 2**, the temperature used here is 30 °C, and thus the LPP cannot be observed by self-assembly without a template. Such template may also be provided in the form of a water layer (**Fig. 3**).

**Figure 2.**
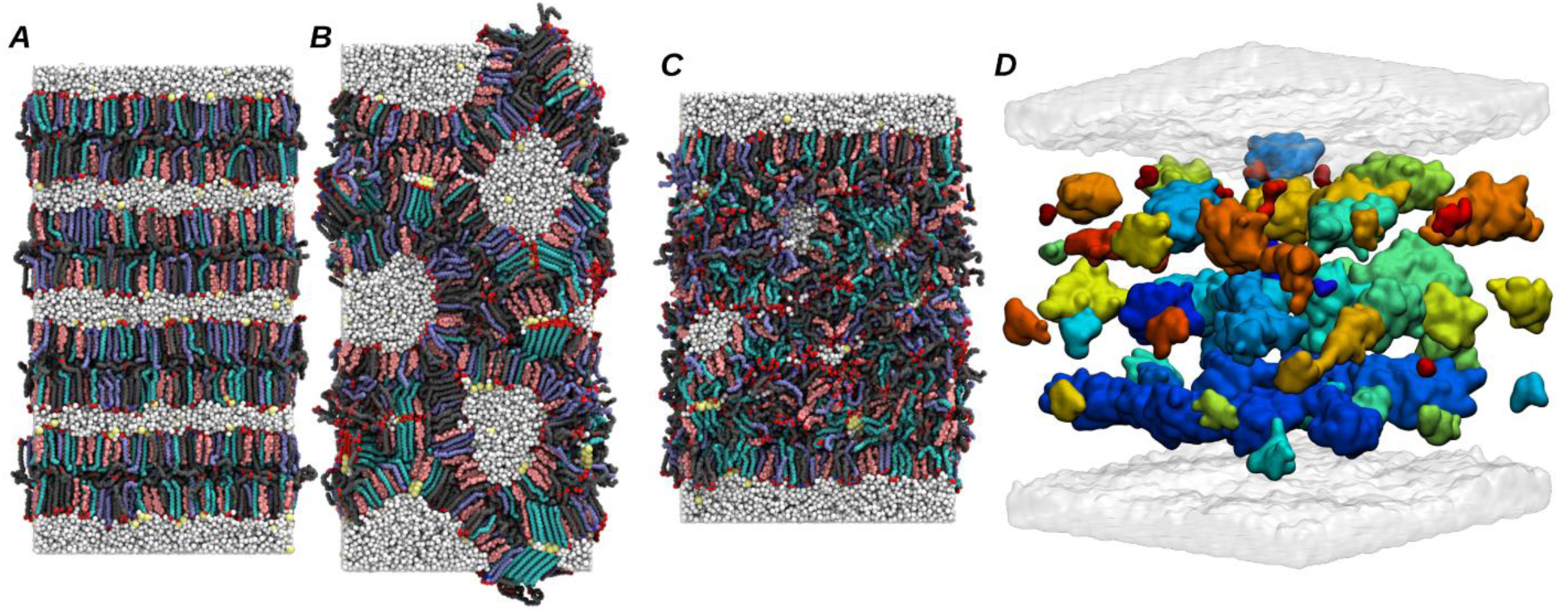
Skin lipid bilayers subject to heat reach a hemi-fused state with interstitial water confined into droplets or channels. Panel **A** shows the initial snapshot of the 5:1 water:lipid 16×16×32 nm^3^ atomistic model, and **B** the same system after heating at 95 °C for 0.2 µs followed by annealing 30 °C for 1.8 µs; small crystalline domains are formed during the latter stage. Panel **C** shows the final snapshot of the 2:1 water:lipid model after heating. Lipid molecules are colored as in **Fig. 1**, water in white and Na^+^ ions in yellow. Panel **D** shows the final snapshot of the 16×16×32 nm^3^ 2:1 water:lipid model, showing the distribution of interstitial water as clusters shown in distinct colors.

**Figure 3.**
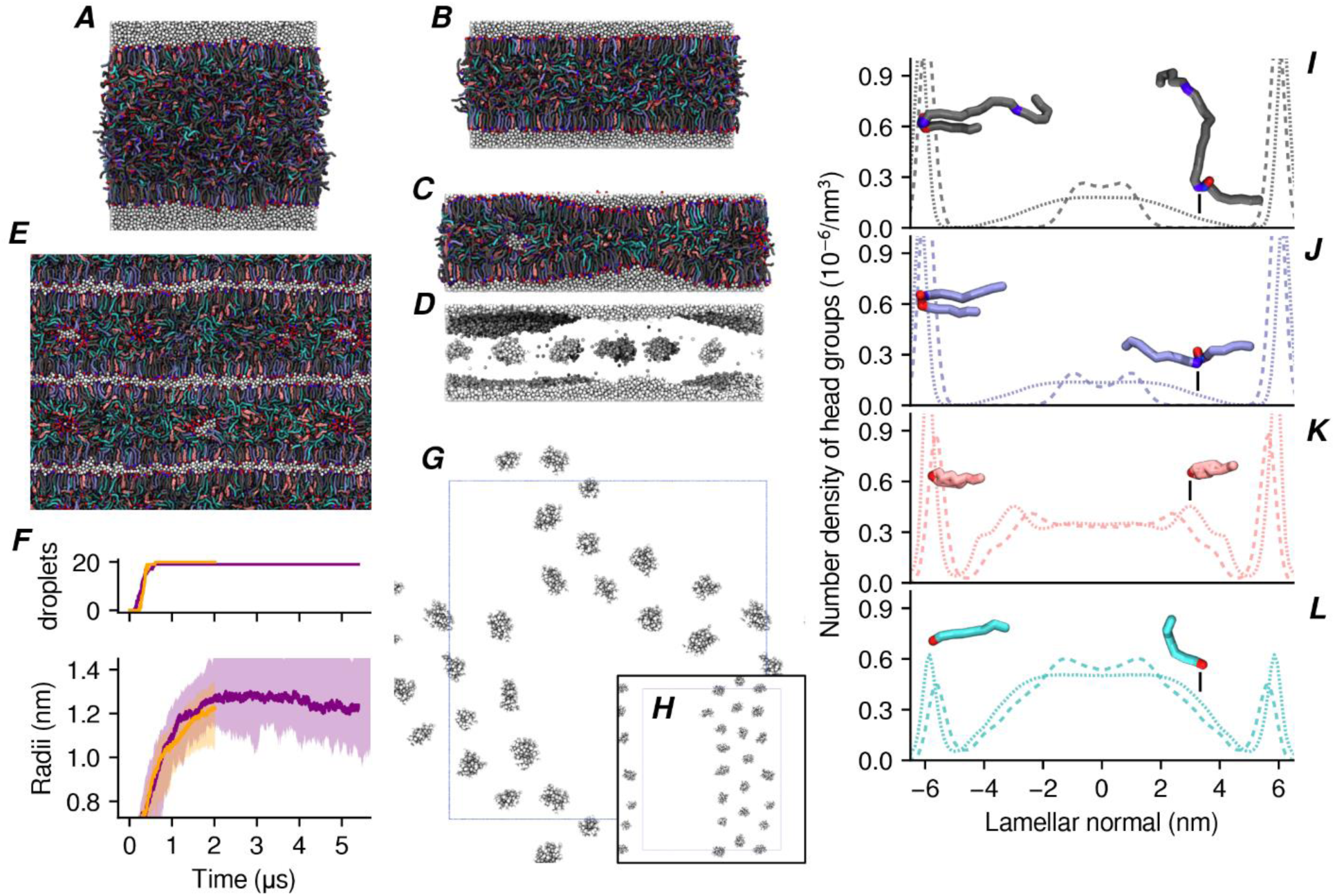
Self-assembled model lamellae with thickness and lipid distribution consistent with the LPP. Initially randomized lipids **(A)** reorganize into dehydrated lamellae with homogeneous thickness **(B)** or hydrated lamellae with inhomogeneous thickness **(C)**; multi-lamellar models are also simulated **(E).** Hydrated lamellae contain water droplets **(D)** formed by nucleation and growth of water molecules; **G** and **H** show the final distributions of droplets (seen from above the lamellar plane) for two independent runs. The number and the median, maximum and minimum of the droplets’ radii are shown in **F** as a function of simulated time. Panels **I-L** show the distribution of lipid head groups for the dehydrated lamella shown in **B** (dotted lines) and the hydrated lamellae shown in **E** (dashed lines).

In mixtures containing cholesterol, saturated chains are packed following a predominantly hexagonal geometry lacking long-range crystalline order (**Fig. S4**), a structure characteristic of the L_o_ lipid phase. Cholesterol molecules may remain aligned with the saturated chains, or position near the isotropic liquid-disordered linoleic-acid segments of CER[EOS]; consistent with neutron diffraction studies, the relative probability of the latter state increases at lower humidity (**Fig. S6**). The boundary between the liquid-ordered outer layers and the liquid-disordered interior as the inflection point of the computed electron-density profile, located at *z* = 1.25 nm above the SPP bilayer’s midplane, a value which also delimits the distribution of the terminal methyl groups of the linear lipids (**Fig. 1E**).

### Permeability calculations of small molecules

The bilayer model discussed in **Fig. 1** was tested by computing potentials of mean force (PMFs) for the permeation of eight molecules that are moderately soluble in lipids (octanol-water partition coefficients *K*_ow_ ranging from 0.2 to 5,000), and one molecule with poor solubility in lipids (mannitol, *K*_ow_ = 8×10^−4^). All have a molecular weight below 300 Da and small anisotropy and internal flexibility, thus being amenable to PMF computations using the center-of-mass position along the bilayer’s normal (see Supplementary Methods). The SPP bilayer is potentially also a transferable permeability model because of its well-defined structure.

The PMFs computed across the hydrated SPP bilayer with atomistic detail span the interval from *z* = 0 (bilayer’s midplane) to *z* = 4 nm (bulk water phase) (**Fig. S8**). Near the lipid-water interface (*z* = 2.7 nm) no deep free-energy minima characteristic of stable adsorption can be observed, due to mild or absent amphiphilicity of the molecules examined, and the tight lipid packing in the L_o_ outer leaflet. For each PMF, the largest free-energy barrier is located near the order-disorder boundary (**Fig. 1E**). It is reasonable to attribute this barrier to the entropic cost of disentangling chain segments around the small molecule, whose movement as it progresses into the L_d_ region is constrained laterally by the saturated lipid chains. This effect is, however, only thermodynamic, because the computed diffusion coefficient is constant across the bilayer and particularly across the order-disorder boundary (**Fig. S9**).

Given the structure of the SC, viable diffusion pathways are not distributed homogeneously in three dimensions but follow the intrinsically two-dimensional lipid matrix. Accordingly, the diffusion path length λ_0_ is known to represent a path that is not straight through the lipid matrix (Mitragotri et al., 2011) and must be accounted for empirically. Unlike in other permeability calculations by MD simulation, the value of λ_0_ is not postulated from simplified models but is obtained by comparing computed and measured data for multiple molecules as follows.

Similarly to the Potts-Guy equation (Potts & Guy, 1992), *k*_P_ is parameterized as a function of the mean diffusion coefficient *D* and of the probability *P*_liq_ that the molecule reaches the more permeable liquid portion of the lipid matrix:

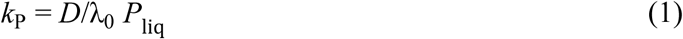

The main difference with the original equation (Potts & Guy, 1992) is that *P*_liq_ measures the solubility of the molecule not in the entire lipid matrix, but only the permeable liquid-disordered portion of it. *P*_liq_ is then estimated by the Boltzmann factor exp(–δG_o/d_), where δG_o/d_ is the free-energy difference between the lowest-lying state in the L_o_ layer and the free-energy barrier associated to the transition into the more permeable L_d_ layer (**Fig. 1E**). The diffusion coefficients *D* computed for several molecules are very similar to the volume-based empirical estimate from the Potts-Guy equation (Potts & Guy, 1992): *D* ∼ exp(–0.0061×MW) where MW is the molecular weight (**Fig. S9**), confirming that steric hindrance is the main factor in determining diffusion for small molecules, and allowing to treat diffusivity and lipophilicity as independent factors (Mitragotri et al., 2011).

A linear regression of (log(*P*_lip_)+log(*D*)) against the experimental log(*k*P) yields a slope of 1.03 (*r*^2^ = 0.89) and an intercept of –0.96, validating the use of **Eq. 1**. Based on these parameters, the theoretical maximum of *k*P is approximately 8.9 cm/h, corresponding to a particle with zero volume and 100% probability to reach the L_d_ region. By comparing the computed diffusion coefficients *D* and those modeled empirically in the Potts-Guy equation (Potts & Guy, 1992), the theoretical diffusion coefficient for the same idealized molecule can be estimated as *D*_0_ = 0.052 cm^2^/h, which gives an estimate of λ0 ≈ 59 µm for the diffusion path length. This value is significantly shorter than what is used in most empirical permeability models (Mitragotri et al., 2011), because the probability of accessing a viable diffusion pathway is not factored implicitly into λ0 but is estimated more directly by *P*_lip_. At the same time, the value of λ0 here obtained exceeds the typical thickness of the SC (∼25 µm) by less than an order of magnitude, consistent with a permeation model following the boundaries between corneocyte cells rather than a straight path.

Although the computed values of *k*_P_ agree well with measured ones for all lipophilic molecules (**Fig. 1F**), the permeability of a strongly hydrophilic species such as mannitol using the above parameters is strongly underestimated (1.6×10^−7^ cm/h vs. 3.7×10^−5^ cm/h). Such discrepancy is in line with similar computations across other model skin lipid systems (Lundborg et al., 2018), and is a known limitation of models based on homogeneous lipid lamellae (Mitragotri et al., 2011). There is in fact an increasing consensus that the permeability of hydrophilic molecules is explained by a distinct pathway from the lamellar phase (Flynn, 1989; Kasting et al., 2019; Menon & Elias, 1997; Mitragotri et al., 2011).

### Bilayers fuse easily at high temperature

While the molecular mechanism of formation of the LPP in the SC is currently unclear, its formation in synthetic models requires heating of the mixture near the range of melting temperatures of the ceramides (Bouwstra et al., 1996). It is therefore useful to identify any meta-stable states produced by heating of skin lipids, particularly in the presence of water: part of this goal can be achieved by simulating the heating of SPP bilayers.

Stacks of four hydrated bilayers were prepared with the same composition discussed above and two model sizes of 16×16×32 nm^3^ and 24×24×32 nm^3^ (**Fig. 2**). For the smaller system size, two distinct pH conditions were modeled using fully protonated FFAs (low pH), and an equal mixture of protonated and unprotonated FFAs (intermediate pH). For both systems, 2:1 as well as 5:1 water:lipid ratios were considered.

Atomistic simulations at 30 °C for approximately 0.5 µs did not reveal significant changes in the multi-bilayer structure. A short heating cycle was then applied to each model system (0.25 µs at 95 °C, well above the melting temperatures of most ceramides) to enhance the rate of structural changes. The large temperature promoted the hemi-fusion between bilayers at multiple contact points (**Fig. 2B**,**C**), and the reorganization into a inverse-hexagonal structure. Mixing of lipids from the fused bilayers further consolidated this structure (**Fig. 2B**).

The hemi-fused lipid regions contain a significantly increased proportion of CER[EOS] in extended form (**Fig. S11**), similarly to the LPP structure. This transition is concurrent with lipid mixing, and resembles a widely used model for the early stages of phospholipid membrane fusion (Stevens et al., 2003). After hemi-fusion, water is organized as continuous channels in the 16×16×32 nm^3^ model at 5:1 water:lipid (**Fig. 2B**), and as a combination of channels and droplets in the 2:1 water:lipid model and the larger 24×24×32 nm^3^ models (**Fig. 2C**,**D**).

After heating, the temperature was brought down to 30 °C to investigate any metastable structures. The resulting structures maintained the overall geometry assumed during heating: however, exact hexagonal spacing in the 16×16×32 nm^3^ models is prevented by the uneven thicknesses of the lipid regions surrounding the water channels. Due to the limited simulation time scale, the hemi-fused regions retain significant double-bilayer structure and do not progress toward the LPP. This configuration was also found to be meta-stable in recent CG simulations of comparable length, where a lateral compression was applied to the lipids to enhance lipid order (Moore et al., 2018): this compression was not used here, and each unit cell vector was pressure-controlled independently at 1 atm.

### Skin lipids self-assemble into lamellae of thickness compatible with the LPP

To model skin lipid lamellae with thickness above the SPP, 20-nm thick lipid slabs were prepared and allowed to reach an equilibrium structure via CG simulation. A water buffer was used to allow anisotropic transformations in the unit cell without artifacts imposed by the periodic boundary conditions. The lipid/water interfaces were initialized based on two hydrated lipid bilayer leaflets (**Fig. 1**), while the inner lipids (a total of ∼15 nm thickness) were set up in randomized conditions (**Fig. 3A**). The initial dimensions of the unit cell are approximately 27×27×26 nm^3^, including the water buffer at 10:1 water:lipid ratio. After equilibration, excess water was removed to generate multi-lamellar stacks (**Fig. 3E**), as discussed below.

Two lipid slabs were simulated at 1 atm using fully anisotropic and semi-anisotropic pressure couplings: no restrictions were placed on the movement of water and these models are referred to as “hydrated” models. One of the two slabs was also simulated with an additional boundary potential applied selectively to the water particles, preventing them from entering the region within ±6 nm from the midplane: this model is hereafter called “dehydrated”, reflecting the lack of water in the lamellar interior regardless of the water buffer.

Lipid molecules progressively diffuse over 0.5 μs from the randomized inner layer to the surface of the lamella, decreasing its thickness and increasing its total surface area by up to 60% (**Fig. 3B**,**C**). Within 2 μs, the unit cell dimensions of each system converge in all models. For the “dehydrated” system, the final lamellar thickness is approximately 13 nm, and is determined by interplay between intermolecular forces and the boundary potential acting on the water particles (**Fig. 3B**).

The two “hydrated” lamellae reach an equilibrium structure with two apparent coexisting thicknesses of ∼11 nm and ∼6 nm, respectively (**Fig. 3C**). In the thick regions, the transformation is accompanied by the aggregation of water molecules into small droplets in the lamellar interior (**Fig. 3D**). While appearing as containing significant water, the droplets themselves account for only 1.7 water molecules for every lipid molecule that is not part of the outer leaflets, and thus represent a level of local humidity compatible with the SC.

Homogeneous lamellae were obtained by extracting an 8×8 nm^2^ section of one of the thick regions and removing all water in excess of 5:1 water:lipid; one droplet was retained in the model, and counted as part of the 5:1 ratio. This section was then equilibrated for 25 μs with periodic boundary conditions and anisotropic pressure coupling, reaching a thickness of about 13 nm (**Fig. 4A**,**C**). The resulting snapshot was also replicated 4×4 times parallel to the plane and 2 times orthogonally (32×37×26 nm^3^ dimensions) and equilibrated for 2 μs, yielding the multi-lamellar structure shown in **Fig. 3E**.

**Figure 4.**
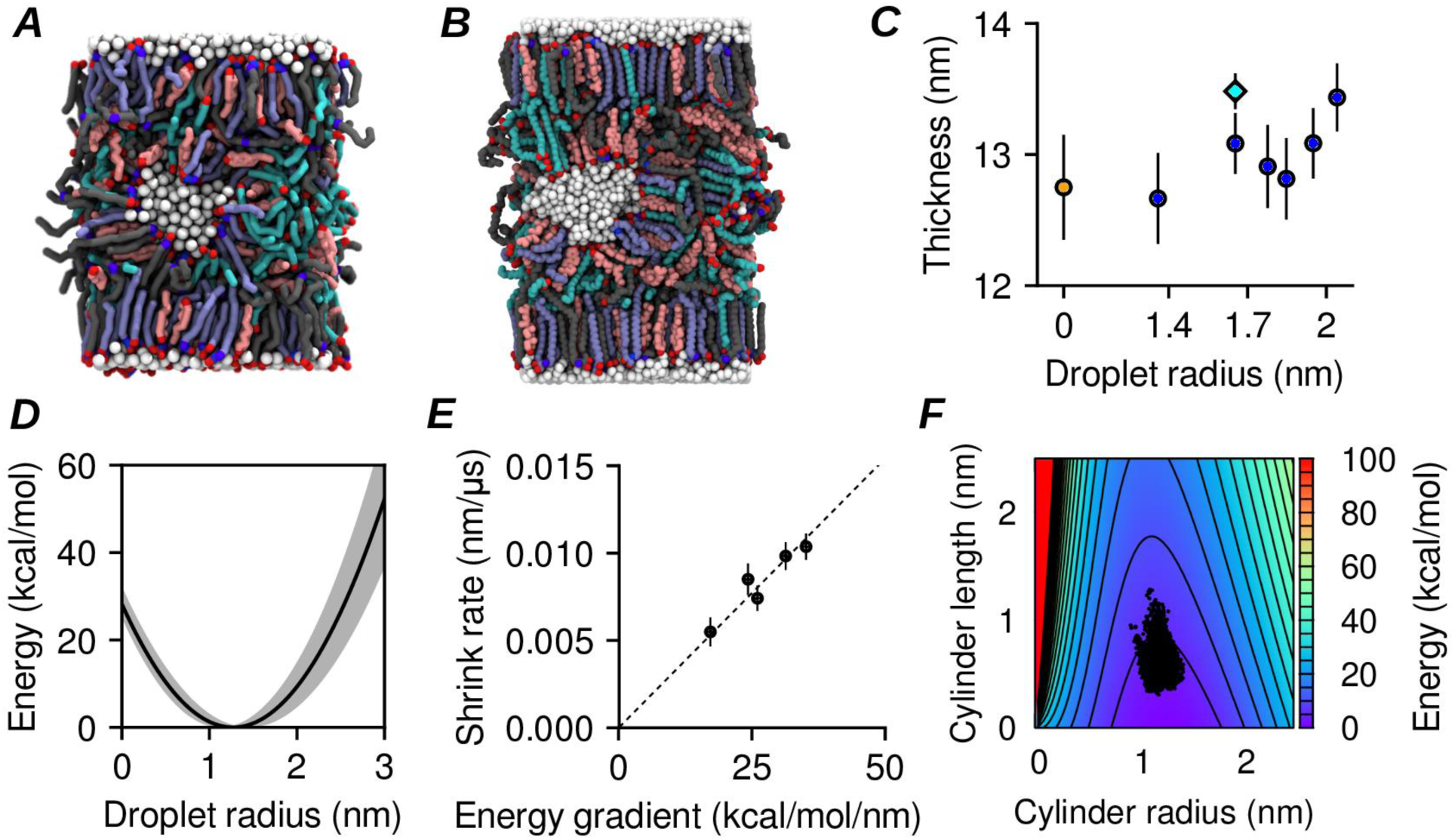
Water droplets are meta-stable within a lamellar core. Panel **A** shows a CG simulation snapshot after 25 µs, and **B** a snapshot from a 2 µs atomistic simulation. **C** shows the estimated thicknesses of the dehydrated CG lamella (orange circle), of the hydrated CG lamellae (blue circles), and of the hydrated atomistic lamella (cyan diamond). **D** shows the energy as a function of the droplet radius from **Eq. 2**; the shaded area reflects the uncertainty in the interfacial tension estimate. **E** shows the estimated rate of shrinkage of the larger droplets as a function of the gradient of **Eq. 2** with respect to the droplet’s radius. **F** shows the energy landscape for a droplet shaped as a cylindrical capsule; dots indicate the radii and lengths distribution for self-assembled droplets (**Fig. 3**). Error bars are standard deviations.

The resulting lipid distribution of the ∼13 nm thick lamellae is symmetric around the midplane, consistent with the structure factors measured by X-ray and neutron diffraction (Groen et al., 2009; Mojumdar et al., 2015a). The interior exhibits a partially de-mixed lipid distribution characteristic of the LPP (Beddoes et al., 2018; Bouwstra et al., 2001; Mojumdar et al., 2015a), with a higher concentration of ceramides in the outer leaflets (±4 to ±6 nm), of cholesterol in the regions immediately below the outer leaflets (±2 to ±4 nm), and of fatty acids in the central region (–2 nm to +2 nm) (**Figs. 3J-L** and **S14**). This de-mixing is caused by the selective migration of lipids during equilibration and was not determined by the initial conditions.

In all models, the computed electron density profile (**Fig. S14)** shows well defined peaks at the outer edges of the lamella and a broader region of high density near its core: the magnitude of the latter depends on the presence of interstitial water (“hydrated” vs. “dehydrated” models). The low-density regions are located approximately at ±4 nm, consistent with X-ray diffraction (Groen et al., 2009). The distribution of lipid headgroups (**Fig. 3J-L**) further reveals that only the outer leaflets are well defined: instead, lipids in the central region are relatively isotropic in the dehydrated lamella, or spherically arranged around the droplets in the hydrated models. This is in contrast with the structural models derived from X-ray and neutron diffraction (Groen et al., 2009; Mojumdar et al., 2015a), which indicate the presence of inner peaks at ±2 nm, less intense than the outer peaks but sufficiently narrow to induce diffraction. It is possible that the difference may be caused by the CG model lacking the capability to form ordered hydrogen-bonds between the head groups, or the CG water particles accounting for three water molecules each and preventing isolated water molecules to act as an intermediate between head groups. Due to this discrepancy, the dehydrated structure shown in **Fig. 3B** is only an approximation to the LPP.

In the hydrated lamellae, the ceramides’ head groups are exposed to liquid water from either the outer layer or the droplet, and the limitations of the CG model discussed above do not apply, suggesting that the corresponding structure is meta-stable. Nonetheless, possible biases of the CG lipid model towards inverse-micellar structures were investigated by constructing hexagonally-packed lattices of droplets and simulating them at CG resolution (**Fig. S15**): this structure was unstable, and within 10 μs evolved into a coexistence of inverse-micellar and lamellar phases.

The final snapshot after 25 µs of a 8×8 nm^2^ periodic lamellae containing a 1.6-nm droplet (**Fig. 4A**) was also used to initialize an atomistic simulation (**Fig. 4B**). Despite significant diffusion of water molecules between the droplet and its periodic images (**Fig. S17**) and of the lipids surrounding the droplet (**Fig. S18**) over 2 µs, the net transfer of water between the droplet and the outer layer is almost zero (**Table 1**), and the droplet does not dissolve within this timescale. Furthermore, the droplet retains its near-spherical shape (**Fig. 4B**) and does not transition to a water layer. Data from this atomistic simulation are also used to compute the electron densities shown in **Fig. S14**. These results indicate that interstitial water droplets in the interior of skin lipid lamellae are relatively long-lived.

**Table 1.** Water transfer statistics for a single lamella with one water droplet and one water layer from CG and atomistic simulations. Rates are computed as described in Supplementary Methods, over the entire atomistic trajectory and a CG trajectory segment where no significant changes in the droplet size were observed (as confirmed by equal transfer rates to and from the water layer). The column headings follow the convention that the first label indicates where the water particle started from, while the second label indicates where it went. Transfers of water molecules between the single droplet and its periodic images, though occurring in both simulations in significant numbers, were excluded from the analysis because of the ambiguity in the length of their travel time.

| | CG Data (~7.5 μs) | | Atomistic Data (~2 μs) | |
| --- | --- | --- | --- | --- |
|  | Layer-Droplet | Droplet-Layer | Layer-Droplet | Droplet-Layer |
| **Number of Transfers** | 2037 | 2058 | 0 | 1 |
| **Exchange Rate (ns<sup>-1</sup>)** | 0.273 | 0.267 | 0 | 0.0008 |
| **Mean Displacement ± Standard Error (nm)** | 6.6 ± 0.1 | 6.4 ± 0.1 | N/A | 4.93 |

### Excess water aggregates into interstitial droplets of controlled size

After nucleation, droplets grow exponentially up to a target radius of *r*^*^ = 1.3 nm (**Fig. 3F**); 19 and 20 droplets are formed in each hydrated lamella, positioned along a quasi-hexagonal two-dimensional lattice (**Fig. 3G**,**H**) and separated from each other by lipid regions of thickness comparable with a lipid bilayer (SPP). No long-range positional order can be observed in the lateral distribution of the droplets.

The size distribution of the self-assembled droplets also agrees well with the lower end of the distribution of droplets or channels resulting from atomistic simulations with heating (**Figs. 2 S19**). Atomistic distributions of droplets above 1.3 nm could not be compared, because they were determined by the initial water content of those atomistic simulations. A model energy function for the droplets’ size was thus defined as follows. The free-energy of the surface *S* of a water droplet was modeled as the sum of the interfacial tension with the lipid phase and the elastic bending energy of the surrounding lipid layer (Helfrich, 1973):

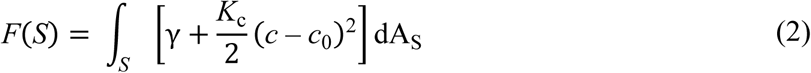

where γ is the tension of the water/lipid interface, *c* = *r*_*x*_^−1^*+r*_*y*_^−1^ and *c*_0_= *r*_0_^−1^ are the total and spontaneous curvatures of the lipids, *K*_c_ is their bending modulus, and dAS is the surface area element. Lacking specific measurements, the interfacial tension between water and 1-octanol (γ ≈ 8.5 mN/m) was used, and for simplicity assumed not to change with the curvature *c*. Based on the distribution of interfacial tension values of fatty alcohols, 2 mN/m was used as the uncertainty (**Fig. 4C**).

The radius of spontaneous curvature *r*0 could be obtained at conditions of high local humidity, where all lipid head groups are exposed to water and the interfacial tension term in **Eq. 2** is constant. Currently, no direct measurements of *r*0 are available. Based on the permeabilities of many hydrophilic species across skin samples a probability distribution of the radii of water channels through the SC has previously been derived *P*(*r*) ∼ exp(–0.045×*r*^2^). The mean of this distribution is approximately 2.7 nm (Tezel et al., 2003). This number is also compatible with the range of the radii of ceramide-lined channels in phospholipid membranes, 0.8 to 11 nm (Siskind et al., 2002), and was used here as the radius of spontaneous curvature *r*_0_. To estimate *K*_*c*_ for the skin lipid mixture, the computed area compressibility modulus of the SPP bilayer (*K*_*A*_ = 273 ± 35 mN/m) was used with the polymer-brush model (Rawicz et al., 2000) to obtain *K*_*c*_ = 9.5 ± 1.2 kcal/mol.

For a spherical droplet, the free-energy estimated by **Eq. 2** has a minimum at a radius of approximately *r*^*^ = 1.3 nm (**Fig. 4C**), consistent with the target radius of the self-assembled droplets (**Fig. 3F**). **Eq. 2** was then also tested by constructing additional copies of the single-droplet model (**Fig. 4A**,**B**), each with an increased number of water particles up to 6 times the original droplet. After minimization and equilibration with semi-anisotropic pressure coupling, all of the larger droplets gradually lose water, shrinking with an approximately exponential decay (**Fig. S20**). The rates of shrinkage are proportional to the free-energy gradient d*F*(*r*)/d*r* calculated at the corresponding radii (**Fig. 4E**), confirming the validity of the model to describe the dynamics of the droplets.

The decay time constants range between approximately 75 and 100 µs (**Fig. 4E**) with a mean of approximately 90 µs, much longer than the initial nucleation and growth of the droplets (**Fig. 3F**). Because the exchange rates of individual water particles are relatively high in the CG model (**Table 2**), such long decay times are not water-limited. Rather, lipid movements control droplet decay times given that the surface area of a droplet, and hence its size, is limited by the movement of the lipid head groups surrounding it.

**Table 2.**
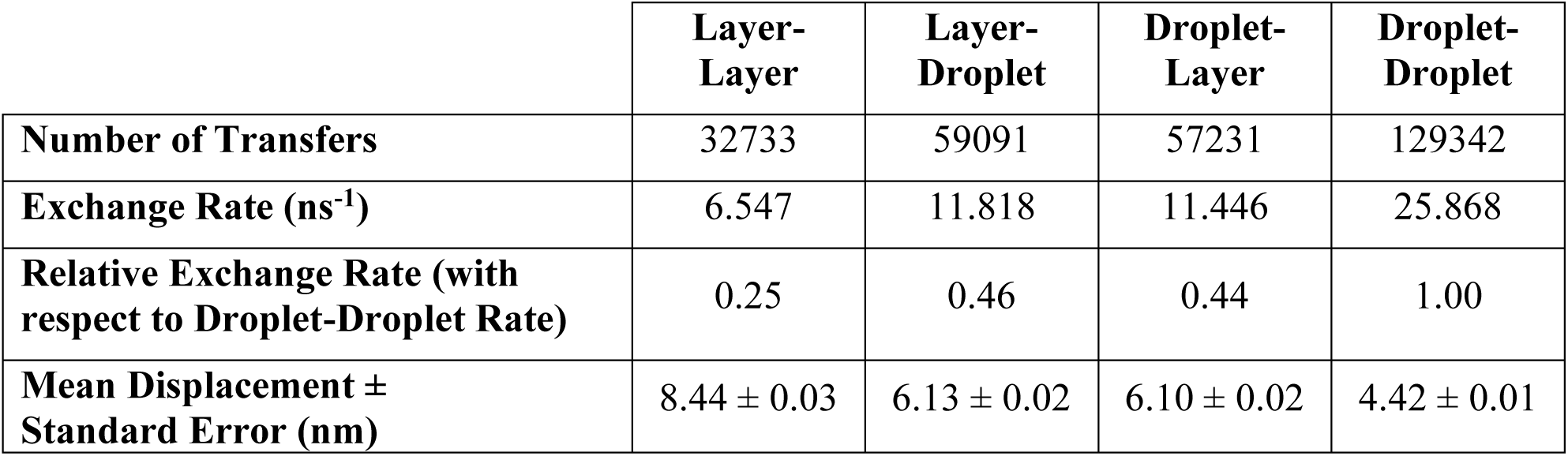
Water exchange events in a system with multiple droplets at CG resolution. These results are based on 5 μs of CG simulation with two lamellae and 38 droplets initially (shown in **Fig. 5A**) and were determined using the same approach as for **Table 1**. Due to limited sampling of merging events, it is not possible to compute directly the net water transfer rates between individual droplets. Based on a decay rate of 90 µs evaluated with the single-droplet system (**Fig. 4E**) and the 45% ratio between droplet-layer and droplet-droplet water exchange rates estimated here, a decay time of 40 µs is estimated for droplet-droplet net water transfer. Because diffusion times of CG lipids are shorter than atomistic lipids by about an order of magnitude (**Fig. S18**), this value is probably underestimated.

|  | **Layer-Layer** | **Layer-Droplet** | **Droplet-Layer** | **Droplet-Droplet** |
| --- | --- | --- | --- | --- |
| **Number of Transfers** | 32733 | 59091 | 57231 | 129342 |
| **Exchange Rate (ns<sup>-1</sup>)** | 6.547 | 11.818 | 11.446 | 25.868 |
| **Relative Exchange Rate (with respect to Droplet-Droplet Rate)** | 0.25 | 0.46 | 0.44 | 1.00 |
| **Mean Displacement ± Standard Error (nm)** | 8.44 ± 0.03 | 6.13 ± 0.02 | 6.10 ± 0.02 | 4.42 ± 0.01 |

### Elongation of water droplets into channels is more favorable than isotropic growth

The above data indicate that the formation of continuous water channels is a rare event, which may be promoted by external factors. To estimate the energy cost of such an event, the free-energy of a model cylindrical capsule was computed using the model from **Eq. 2** (**Figs. 4F** and **S21**). Using the leading eigenvalue of each droplet’s inertia tensor to parameterize the length *L*, and the mean of the other two eigenvalues for the radius *r*, the shape fluctuations of self-assembled droplets were compared to the energy landscape. All droplets are distributed near the minimum of the free-energy *F*(*r, L*), located at *r*^*^ = 1.3 nm and *L*^*^ = 0 (**Fig. 4F**). Due to thermal fluctuations of the droplet’s shape, the smallest cylindrical length explored is however not zero, but *L*_min_ ≈ 0.3 nm, i.e. the approximate radius of a water molecule. The largest length explored is *L*_max_ ≈ 1.2 nm, corresponding to droplets that are about 50% longer along one axis than the other two axes. The distribution of individual droplets exhibits tails that appear to follow the constant volume contour lines (**Fig. S21**) but remain within 8 kcal/mol from the free-energy minimum. Therefore, the two-dimensional model is a simplified but reasonably accurate description of droplet anisotropy.

Assuming a droplet with optimal initial radius *r* = 1.3 nm, the elongation required to cover the typical distance with the nearest droplets is *L* ≈ 6 nm. If the transformation occurs at constant volume (i.e. a fixed size for the droplet), approximately 43 kcal/mol of energy would be required (**Fig. 4F)**. If changes in the droplet’s volume are allowed by addition of water, this activation energy is reduced to 33 kcal/mol. Both values are sufficiently high to justify a low probability of water channels at equilibrium conditions (Tezel et al., 2003). At the same time, corrugations induced by the local surface of the corneocytes (Hou et al., 1991), as well was the use of trans-dermal delivery enhancement methods, could offset at least part of this energy cost, greatly increasing the probability of formation of water channels.

### The coalescence of water droplets is affected by interactions between lamellae

Although a multi-lamellar model with embedded droplets is meta-stable with homogeneous lamellae and flat equilibrium structure (**Fig. 3E**), it is useful to investigate the dynamics of interstitial water in the presence of corrugations. A simple test of this effect was constructed by replicating one of the snapshots at 2 µs shown in **Fig. 3** along the direction normal to the lamella (**Fig. 5A**). Within 1 μs, the thick regions of the two lamellae move such that the thickness of the water layers becomes homogeneous throughout the unit cell, thus adapting to the irregular lamellar profile (**Fig. 5B**).

**Figure 5.**
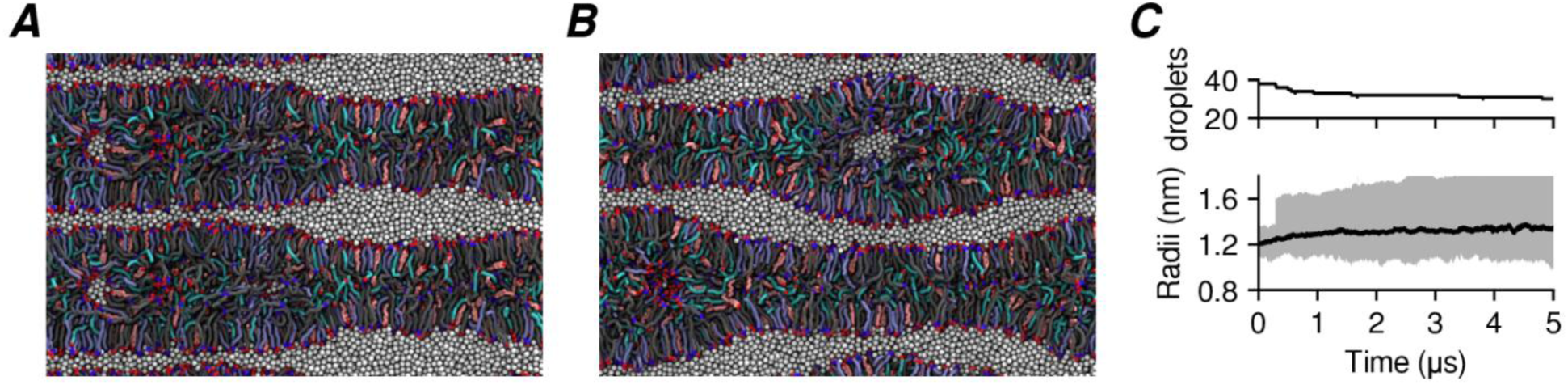
Relative movements of the lipid lamellae influence the number of droplets but not their size distribution. Panel **(A)** shows the initial condition and **(B)** the final snapshot after 5 µs. **(C)** shows the total number of droplets and radii distribution in the two lamellae; the solid line is the median and the shaded area the size range. Note the sudden jump at ∼0.3 μs reflecting the first merging event.

While the thick regions of each lamella maintain their pre-existing thickness, their area along the lamellar plane decreases by about 25-30% (visual inspection). This change is accompanied by the merger of several droplets, starting at 38 droplets in each lamella and reaching 30 droplets after 5 μs (**Fig. 5C**). Despite the decrease in the number of droplets, the total amount of water contained in the droplets increases by ∼15%, and the median droplet radius saturates near 1.3 nm after ripening and merging of individual droplets (**Fig. 5C**).

## DISCUSSION

The results presented here demonstrate a direct connection between the coexistence of multiple lipid phases (**Fig. 6**) and the ability of skin lipids to sustain interstitial water, which is essential in controlling the skin’s permeability to water-soluble substances. Among the phases investigated are bilayers (**Fig. 1**), inverse-hexagonal and inverse-micellar structures (**Fig. 2**), and lamellae with thickness equal to the LPP (**Fig. 3**). A lamella without interstitial water self-assembled in CG simulations has a similar structure to the LPP, with two notable differences: firstly, significant disorder remains in the inner lipids, with intra-lamellar “peaks” being about 3 times broader than in X-ray and neutron diffraction-based models (Groen et al., 2009; Mojumdar et al., 2015a). Secondly, the outer leaflets of the lamella do not appear to show a preference for the long-chain CER[EOS] vs. the shorter-chain CER[NS].

**Figure 6.**
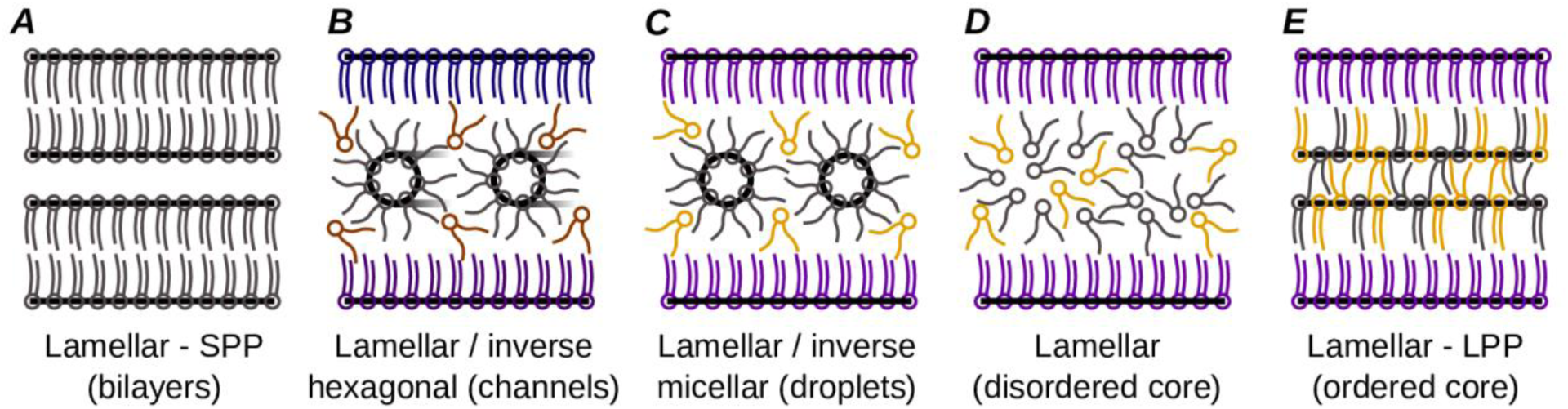
Summary of the stable and meta-stable skin lipid phases relevant to interstitial water. The structures simulated in this study (**A-D**) are compared to the LPP **(E)**: in each panel, ordered interfaces are highlighted with black lines. Lipids are colored in gray if their local composition matches the global composition, purple when enriched in ceramides, and orange when depleted in ceramides. Transitions from bilayers (A) to inverse-hexagonal and inverse-micellar phases (B,C) were observed as a result of heat (**Fig. 2**). Thick lamellae were also obtained by self-assembly (**Fig. 3**) in the hydrated state (C) and dehydrated state (D), with partial transitions between the two as a result of water diffusion. Permeability of hydrophilic molecules through the skin is thought to be non-vanishing only in the presence of meta-stable water channels (B).

Because self-assembly simulations of skin lipids cannot be carried out at atomistic detail, it is difficult to attribute these two discrepancies to the CG force field. Recent atomistic simulations (Wang & Klauda, in press) initialized with the “sandwich” LPP model (Beddoes et al., 2018; Bouwstra et al., 2001; Mojumdar et al., 2015a) show that this model is meta-stable on the µs time scale at 5:1 water:lipid, but unstable at 2:1 water:lipid. It is unclear if this result is due to experimental uncertainty in the water content, or the inability of simulation models to sustain a dehydrated phase of the skin lipids. Due to the use of a pre-initialized structure, the thermodynamic stability of the “sandwich” LPP model remains an open question, and it is possible that CG models based on machine learning of atomistic simulations (Moore et al., 2018) could help elucidate the matter.

In hydrated lamellae, the spontaneous formation of water droplets (**Fig. 3**) allows for the lipid head groups to interact with water, a physical state that is well described by the CG force field used. Atomistic simulations confirm that lamellae containing droplets are meta-stable over µs timescales (**Fig. 4**), raising the possibility that they could be measured given suitable experimental conditions. Unfortunately, the two-dimensional arrangement of the simulated droplets exhibits locally hexagonal packing but lacks long-range order (**Fig. 3**). Comparison with X-ray diffraction data of a 3D inverse-micellar phospholipid phase (Shearman et al., 2009) suggests that even for a perfect lattice the inverse-micellar peaks may not be intense enough to be detected in X-ray or neutron diffraction.

High resolution cryo-electron microscopy of extracted skin sections shows in some images series of intra-lamellar spots of the same size and spacing of the simulated droplets, with the majority of images showing a purely lamellar structure (Iwai et al., 2012): this supports the hypothesis that inverse-micellar droplets are a meta-stable state of the skin’s lamellae. Beyond structural data, the permeabilities of hydrophilic substances have already been used to model the distribution of radii of water channels (Tezel et al., 2003), and such radii describe quantitatively the self-assembled water droplets as well (**Eq. 2**). Coalescence of water droplets is then estimated to be a rare event (**Fig. 4F**) whose probability can be increased at certain conditions (**Fig. 5**), promoting the formation of hydrophilic permeation pathways.

Besides hydrophilic permeability, another possible function of a pseudo-lattice of droplets inside each lamella is facilitating the relative movement of the two lamellar outer leaflets under shear. This property would be most important before and during maturation of the deeper and more humid layers of the epidermis, and it is likely that droplets would vanish at the lowest humidity levels, leaving a lamella with negligible interstitial water and LPP structure (**Fig. 6**). Observing such transition exceeds available simulation time scales. However, the slower rate of water exchange across the lamellae than between droplets (**Table 2**) supports the hypothesis that the water layers between lamellae evaporate first, while the more stable droplets within them evaporate at a later stage. Consistent with this hypothesis is also the multi-state mechanism of skin homeostasis proposed from studies of evaporation from amphiphiles/water mixtures (Roger et al., 2016). The steep increase in water permeability of the SC above >80% relative humidity (Kasting et al., 2003) can also be explained by the formation of continuous channels that are further stabilized by the resulting swelling of the corneocytes’ keratin mesh (Kasting et al., 2003).

Other factors that may promote the formation of water channels are mechanical forces. For example, the 20 kHz frequency of the ultrasound vibrations used to enhance hydrophilic permeability (Mitragotri et al., 1995) is at the order of magnitude of net droplet-droplet water transfer (τ ≈ 40 µs, **Table 2**). The previously suggested activation energy of 43 kcal/mol per droplet gives an upper bound of about ∼1.9×10^−4^ J/cm^2^ of energy density (assuming 15 lipid lamellae per corneocyte and 15 corneocytes across the SC). The energy density supplied by permeability-enhancing ultrasound vibrations over the course of several hours is >100 J/cm^2^ on the entire epidermis (Mitragotri et al., 1995), of which >3 J/cm^2^ is apportioned to the SC’s lipid matrix based on its volume. Although much of this energy would be lost by thermal dissipation, its total is more than 50,000 times higher than the activation energy required to overcome the energy barrier. This comparison further supports the coalescence between water droplets as a mechanism for hydrophilic permeability.

In conclusion, the results here presented demonstrate that the multiplicity of phases of the skin lipids, rather than being an artifact of different experimental conditions, explains directly the presence of hydrophilic pathways of permeation. The skin’s homeostasis and the modulation of barrier function can thus be described by the relative abundance of stable and meta-stable structures, of which only the latter would provide continuous water channels through the skin.

## METHODS

### Molecular dynamics simulation force-fields for skin lipids

Forces between skin lipids in molecular dynamics (MD) simulations were computed with atomistic and coarse-grained (CG) levels of resolution. Atomistic simulation parameters were taken from the CHARMM36 force field (Klauda et al., 2010; Venable et al., 2014) for all lipids, CGENFF (Vanommeslaeghe et al., 2010) for small molecules and TIP3P (Jorgensen et al., 1983) for water. CG force field parameters for acyl chains, ester groups, hydroxyl groups, cholesterol and water are from published work (MacDermaid et al., 2015; Shinoda et al., 2010). Parameters for amides and the protonated carboxyl group were also derived from experimental properties of the corresponding liquids (see Supplementary Methods).

### MD simulation and PMF calculation protocols

The programs NAMD (Phillips et al., 2005) and LAMMPS (Plimpton, 1995) were employed to carry out MD simulations at the atomic and CG resolutions, respectively. The protocols to treat non-bonded interactions and pressure and temperature coupling are as previously used (MacDermaid et al., 2015) when treating atomistic and CG bilayers. Unless otherwise noted, simulations were carried out at a target temperature of 30 °C and pressure of 1 atm; integration steps were 2 fs and 10 fs for atomistic and CG simulations, respectively. For each simulation, the pressure coupling was isotropic (*NPT* ensemble) until the density of the system reached equilibrium (typically less than 1 ns), and anisotropic (*NP*_*x*_*P*_*y*_*P*_*z*_*T* ensemble) thereafter. Simulations with semi-anisotropic pressure coupling (*P*_*y*_ and *P*_*z*_ coupled) are indicated in the Results. Adaptive biasing force (ABF) simulations (Darve & Pohorille, 2001) were carried out on multiple overlapping windows spaced by 0.1 nm. Further details are in the Supplementary Methods.

### Preparation of initial structures for atomistic and CG simulations

Unless otherwise noted in the Results, CG simulations were initialized as fully randomized mixtures of lipids and water. Atomistic bilayers were prepared by assembling a monolayer system containing 32 molecules each of CER[NS], cholesterol, and behenic acid. The fatty acid chains of 16 CER[NS] molecules were extended to into CER[EOS] molecules, with the linoleic-acid tails initially lying parallel to the monolayer’s plane. Each monolayer was replicated and rotated by 180° to give a symmetric bilayer, which was then solvated with water and neutralized by the addition of Na^+^ ions in the water phase in systems that contained unprotonated FFAs. The resulting 16,000-atoms model (4×4 nm^2^ bilayer area) was then equilibrated and replicated multiple times along the bilayer’s midplane and four times orthogonal to the bilayer. An additional 1.5 µs simulation of the hydrated bilayer was carried out to verify its stability at atomistic detail and mutual agreement with the self-assembled CG bilayers (**Fig. S5**). For PMF calculations, small molecules were inserted in a pre-equilibrated bilayer at different depths and the surrounding lipids re-equilibrated before sampling, to minimize the emergence of spurious barriers associated to non-equilibrium work (Del Regno & Notman, 2018; Gupta et al., 2016). Conversion of CG models to atomistic resolution was done by placing atoms at randomized positions within the radius of the respective CG particle, followed by 20 ns of equilibration at constant volume.

## ACKNOWLEDGMENTS

We are grateful to Grace Brannigan, Richard Pastor, Jeffery Klauda and Samir Mitragotri for useful discussions. G.F. and C.M.M. acknowledge support from the National Science Foundation (NSF) through grants CHE-1212416 and DMR-1120901. Calculations were carried out on the Oak Ridge Leadership Computing Facility Titan supercomputer (INCITE allocation CHM045), and the Temple University Owl’s Nest supercomputer (US Army Research Laboratory contract number W911NF-16-2-0189, NSF major instrumentation grant number 1625061).

## REFERENCES

Beddoes, C. M., Gooris, G. S., & Bouwstra, J. A. (2018). Preferential arrangement of lipids in the long-periodicity phase of a stratum corneum matrix model. Journal of Lipid Research, 59(12), 2329–2338. Doi:10.1194/jlr.M087106

Bouwstra, J. A., Gooris, G. S., Dubbelaar, F. E. R., & Ponec, M. (2001). Phase behavior of lipid mixtures based on human ceramides: coexistence of crystalline and liquid phases. Journal of Lipid Research, 42(11), 1759–1770.

Bouwstra, J. A., Gorris, G. S., Cheng, K., Weerheim, A., Bras, W., & Ponec, M. (1996). Phase behavior of isolated skin lipids. Journal of Lipid Research, 37(5), 999–1011.

Caspers, P. J., Lucassen, G. W., Carter, E. A., Bruining, H. A., & Puppels, G. J. (2001). In vivo confocal Raman microspectroscopy of the skin: Noninvasive determination of molecular concentration profiles. Journal of Investigative Dermatology, 116(3), 434–442. Doi:10.1046/j.1523-1747.2001.01258.

Darve, E., & Pohorille, A. (2001). Calculating free energies using average force. The Journal of Chemical Physics, 115(20), 9169–9183.

Das, C., Noro, M. G., & Olmsted, P. D. (2013). Lamellar and inverse micellar structures of skin lipids: effect of templating. Physical Review Letters, 111(14), 148101.

Del Regno, A., & Notman, R. (2018). Permeation pathways through lateral domains in model membranes of skin lipids. Physical Chemistry Chemical Physics, 20(4), 2162–2174.

Elias, P. M. (2005). Stratum corneum defensive functions: An integrated view. Journal of Investigative Dermatology, 125(2), 183–200. Doi:10.1111/j.0022-202X.2005.23668.

Flynn, G. L. (1989). Mechanism of percutaneous absorption from physicochemical evidence. Percutaneous absorption, 27–51.

Groen, D., Gooris, G. S., Barlow, D. J., Lawrence, M. J., van Mechelen, J. B., Demé, B., & Bouwstra, J. a. (2011). Disposition of ceramide in model lipid membranes determined by neutron diffraction. Biophysical journal, 100(6), 1481–1489.

Groen, D., Gooris, G. S., & Bouwstra, J. A. (2009). New Insights into the Stratum Corneum Lipid Organization by X-Ray Diffraction Analysis. Biophysical Journal, 97(8), 2242–2249. Doi:10.1016/j.bpj.2009.07.040

Groen, D., Gooris, G. S., & Bouwstra, J. A. (2010). Model Membranes Prepared with Ceramide EOS, Cholesterol and Free Fatty Acids Form a Unique Lamellar Phase. Langmuir, 26(6), 4168–4175. Doi:10.1021/la9047038

Gupta, R., Sridhar, D., & Rai, B. (2016). Molecular dynamics simulation study of permeation of molecules through skin lipid bilayer. The Journal of Physical Chemistry B, 120(34), 8987–8996.

Helfrich, W. (1973). Elastic Properties of Lipid Bilayers - Theory and Possible Experiments. Zeitschrift Fur Naturforschung C-a Journal of Biosciences, C 28(11-1), 693–703.

Hirvonen, J., Kalia, Y. N., & Guy, R. H. (1996). Transdermal delivery of peptides by iontophoresis. Nature Biotechnology, 14(13), 1710.

Hou, S. Y. E., Mitra, A. K., White, S. H., Menon, G. K., Ghadially, R., & Elias, P. M. (1991). Membrane structures in normal and essential fatty-acid deficient stratum-corneum - characterization by ruthenium tetroxide staining and X-ray diffraction. Journal of Investigative Dermatology, 96(2), 215–223.

Iwai, I., Han, H., Hollander, L. D., Svensson, S., Ofverstedt, L.-G., Anwar, J., Brewer, J., Bloksgaard, M., Laloeuf, A., Nosek, D., Masich, S., Bagatolli, L. a., Skoglund, U., & Norlén, L. (2012). The Human Skin Barrier Is Organized as Stacked Bilayers of Fully Extended Ceramides with Cholesterol Molecules Associated with the Ceramide Sphingoid Moiety. The Journal of Investigative Dermatology, 1–11.

Jorgensen, W. L., Chandrasekhar, J., Madura, J. D., Impey, R. W., & Klein, M. L. (1983). Comparison of simple potential functions for simulating liquid water. Journal of Chemical Physics, 79(2), 926–935. Doi:10.1063/1.445869

Kasting, G. B., Barai, N. D., Wang, T. F., & Nitsche, J. M. (2003). Mobility of water in human stratum corneum. Journal of Pharmaceutical Sciences, 92(11), 2326–2340. Doi:Doi 10.1002/Jps.10483

Kasting, G. B., Miller, M. A., LaCount, T. D., & Jaworska, J. (2019). A composite model for the transport of hydrophilic and lipophilic compounds across the skin: steady-state behavior. Journal of pharmaceutical sciences, 108(1), 337–349.

Klauda, J. B., Venable, R. M., Freites, J. A., O’Connor, J. W., Tobias, D. J., Mondragon-Ramirez, C., Vorobyov, I., MacKerell, A. D., & Pastor, R. W. (2010). Update of the CHARMM All-Atom Additive Force Field for Lipids: Validation on Six Lipid Types. Journal of Physical Chemistry B, 114(23), 7830–7843. Doi:Doi 10.1021/Jp101759q

Lundborg, M., Wennberg, C. L., Narangifard, A., Lindahl, E., & Norlén, L. (2018). Predicting drug permeability through skin using molecular dynamics simulation. Journal of controlled release, 283, 269–279. Doi:10.1016/j.jconrel.2018.05.026

MacDermaid, C. M., Kashyap, H. K., DeVane, R. H., Shinoda, W., Klauda, J. B., Klein, M. L., & Fiorin, G. (2015). Molecular dynamics simulations of cholesterol-rich membranes using a coarse-grained force field for cyclic alkanes. Journal of Chemical Physics, 143(24), 243144. doi:10.1063/1.4937153

McIntosh, T. J., Stewart, M. E., & Downing, D. T. (1996). X-ray diffraction analysis of isolated skin lipids: reconstitution of intercellular lipid domains. Biochemistry, 35(12), 3649–3653. doi:10.1021/bi952762q

Menon, G. K., & Elias, P. M. (1997). Morphologic Basis for a Pore-Pathway in Mammalian Stratum Corneum. Skin Pharmacology and Physiology, 10(5-6), 235–246. doi:10.1159/000211511

Mitragotri, S., Anissimov, Y. G., Bunge, A. L., Frasch, H. F., Guy, R. H., Hadgraft, J., Kasting, G. B., Lane, M. E., & Roberts, M. S. (2011). Mathematical models of skin permeability: an overview. International journal of pharmaceutics, 418(1), 115–129.

Mitragotri, S., Blankschtein, D., & Langer, R. (1995). Ultrasound-mediated transdermal protein delivery. Science, 269(5225), 850–853.

Mojumdar, E. H., Gooris, G. S., Barlow, D. J., Lawrence, M. J., Deme, B., & Bouwstra, J. A. (2015a). Skin lipids: localization of ceramide and fatty acid in the unit cell of the long periodicity phase. Biophysical Journal, 108(11), 2670–2679. Doi:10.1016/j.bpj.2015.04.030

Mojumdar, E. H., Gooris, G. S., & Bouwstra, J. A. (2015b). Phase behavior of skin lipid mixtures: the effect of cholesterol on lipid organization. Soft Matter, 11(21), 4326–4336. Doi:10.1039/c4sm02786h

Moore, D. J., Rerek, M. E., & Mendelsohn, R. (1997). FTIR spectroscopy studies of the conformational order and phase behavior of ceramides. The Journal of Physical Chemistry B, 101(44), 8933–8940.

Moore, T. C., Iacovella, C. R., Leonhard, A. C., Bunge, A. L., & McCabe, C. (2018). Molecular dynamics simulations of stratum corneum lipid mixtures: A multiscale perspective. Biochemical and biophysical research communications, 498(2), 313–318.

Norlen, L. (2001). Skin barrier formation: The membrane folding model. Journal of Investigative Dermatology, 117(4), 823–829. Doi:10.1046/j.0022-202x.2001.01445

Pham, Q. D., Mojumdar, E. H., Gooris, G. S., Bouwstra, J. A., Sparr, E., & Topgaard, D. (2018). Solid and fluid segments within the same molecule of stratum corneum ceramide lipid. Quarterly Reviews of Biophysics, 51, e7. Doi:10.1017/S0033583518000069

Pham, Q. D., Topgaard, D., & Sparr, E. (2017). Tracking solvents in the skin through atomically resolved measurements of molecular mobility in intact stratum corneum. Proceedings of the National Academy of Sciences, 114(2), E112–E121. Doi:10.1073/pnas.1608739114

Phillips, J. C., Braun, R., Wang, W., Gumbart, J., Tajkhorshid, E., Villa, E., Chipot, C., Skeel, R. D., Kale, L., & Schulten, K. (2005). Scalable molecular dynamics with NAMD. Journal of Computational Chemistry, 26(16), 1781–1802. Doi:Doi 10.1002/Jcc.20289

Plimpton, S. (1995). Fast Parallel Algorithms for Short-Range Molecular-Dynamics. Journal of Computational Physics, 117(1), 1–19. Doi:DOI 10.1006/jcph.1995.1039

Potts, R. O., & Guy, R. H. (1992). Predicting Skin Permeability. Pharmaceutical research, 9, 663–669.

Rawicz, W., Olbrich, K., McIntosh, T., Needham, D., & Evans, E. (2000). Effect of chain length and unsaturation on elasticity of lipid bilayers. Biophysical journal, 79(1), 328–339.

Roger, K., Liebi, M., Heimdal, J., Pham, Q. D., & Sparr, E. (2016). Controlling water evaporation through self-assembly. Proceedings of the National Academy of Sciences, 113(37), 10275–10280. Doi:10.1073/pnas.1604134113

Schmitt, T., Lange, S., Sonnenberger, S., Dobner, B., Demé, B., Langner, A., & Neubert, R. H. (2019). The long periodicity phase (LPP) controversy part I: The influence of a natural-like ratio of the CER [EOS] analogue [EOS]-br in a CER [NP]/[AP] based stratum corneum modelling system: A neutron diffraction study. Biochimica et Biophysica Acta (BBA)-Biomembranes, 1861(1), 306–315.

Schroter, A., Kessner, D., Kiselev, M. a., Hauss, T., Dante, S., & Neubert, R. H. H. (2009). Basic nanostructure of stratum corneum lipid matrices based on ceramides [EOS] and [AP]: a neutron diffraction study. Biophysical journal, 97(4), 1104–1114.

Shearman, G. C., Tyler, A. I., Brooks, N. J., Templer, R. H., Ces, O., Law, R. V., & Seddon, J. M. (2009). A 3-D hexagonal inverse micellar lyotropic phase. Journal of the American Chemical Society, 131(5), 1678–1679.

Shinoda, W., DeVane, R., & Klein, M. L. (2010). Zwitterionic lipid assemblies: molecular dynamics studies of monolayers, bilayers, and vesicles using a new coarse grain force field. The Journal of Physical Chemistry B, 114(20), 6836–6849.

Siskind, L. J., Kolesnick, R. N., & Colombini, M. (2002). Ceramide channels increase the permeability of the mitochondrial outer membrane to small proteins. Journal of Biological Chemistry, 277(30), 26796–26803.

Skolova, B., Janusova, B., Zbytovska, J., Gooris, G., Bouwstra, J., Slepicka, P., Berka, P., Roh, J., Palat, K., Hrabalek, A., & Vavrova, K. (2013). Ceramides in the Skin Lipid Membranes: Length Matters. Langmuir, 29(50), 15624–15633. Doi:Doi 10.1021/La4037474

Stevens, M. J., Hoh, J. H., & Woolf, T. B. (2003). Insights into the molecular mechanism of membrane fusion from simulation: evidence for the association of splayed tails. Phys Rev Lett, 91(18), 188102.

Tezel, A., Sens, A., & Mitragotri, S. (2003). Description of transdermal transport of hydrophilic solutes during low-frequency sonophoresis based on a modified porous pathway model. Journal of Pharmaceutical Sciences, 92(2), 381–393. Doi:10.1002/jps.10299

Vanommeslaeghe, K., Hatcher, E., Acharya, C., Kundu, S., Zhong, S., Shim, J., Darian, E., Guvench, O., Lopes, P., Vorobyov, I., & MacKerell, A. D. (2010). CHARMM General Force Field: A Force Field for Drug-Like Molecules Compatible with the CHARMM All-Atom Additive Biological Force Fields. Journal of Computational Chemistry, 31(4), 671–690. Doi:Doi 10.1002/Jcc.21367

Veiga, M. P., Arrondo, J. L. R., Goni, F. M., & Alonso, A. (1999). Ceramides in phospholipid membranes: effects on bilayer stability and transition to nonlamellar phases. Biophysical journal, 76(1), 342–350.

Venable, R. M., Sodt, A. J., Rogaski, B., Rui, H., Hatcher, E., MacKerell, A. D., Jr., Pastor, R. W., & Klauda, J. B. (2014). CHARMM All-Atom Additive Force Field for Sphingomyelin: Elucidation of Hydrogen Bonding and of Positive Curvature. Biophysical Journal, 107(1), 134–145. doi:10.1016/j.bpj.2014.05.034

Wang, E., & Klauda, J. B. (in press). Molecular Structure of the Long Periodicity Phase in the Stratum Corneum. Journal of the American Chemical Society. doi:10.1021/jacs.9b08995

Wertz, P., Downing, D., & Goldsmith, L. (1991). Epidermal lipids. In Physiology, biochemistry, and molecular biology of the skin (pp.205–236).

White, S. H., Mirejovsky, D., & King, G. I. (1988). Structure of lamellar lipid domains and corneocyte envelopes of murine stratum corneum. An X-ray diffraction study. Biochemistry, 27(10), 3725–3732.

