## Supplementary Material for "Coexistence of lipid phases stabilizes interstitial water in the outer layer of mammalian skin" (from another version of this preprint: version 2, 2019-10-08)

### SUPPLEMENTARY METHODS

#### Coarse-grained (CG) force field parameterization.

CG simulation parameters for the lipids were derived as part this work using
9-6 Lennard-Jones (LJ) with parameters derived using the SDK method (Shinoda et al., 2007). Previous parameters from liquid hydrocarbons, alcohols and lipids (DeVane et al., 2009; Shinoda et al., 2007) were used as-is, and additional parameters were derived from thermodynamic data for formamide ($NH_2CHO$), N-methyl-formamide ($CH_3NHCHO$) and butyric acid ($CH_3(CH_2)_2COOH$). These model compounds were used to represent the head groups of ceramides and free fatty acids, respectively. The same parameters for amide groups were also used in a recent paper to characterize the structure of a liquid crystalline polymer solution (Mogurampelly et al., 2019).

Due to the absence of a dipole moment in the water model that treats three molecules as a spherical particle (Shinoda et al., 2007), and the low water content of the SC's lipid matrix, charged head groups such as $COO^-$ were not modeled in CG simulations, which focus on low pH conditions exclusively.

The following table lists all non-bonded intermolecular potential parameters derived from experimental properties, such as density and surface tension of the reference small molecules. Interactions with water were derived by fitting experimental hydration free-energies. Interactions between pairs of different types (i.e. "non-diagonal" interactions) not listed here were parameterized using a geometric combination rule for the corresponding "diagonal" interaction parameters. The sites are labeled as follows: N = amide NH group, O = carbonyl C=O group, W

= water particle corresponding to three water molecules (Shinoda et al., 2007), COOH = COOH group.

| **CG particle type 1** | **CG particle type 2** | **LJ ε (kcal/mol)** | **LJ σ (Å)** |
|---|---|---|---|
| N | N | 0.2430 | 4.0506 |
| O | O | 0.3233 | 3.7880 |
| N | O | 0.5393, | 3.6246 |
| N | W | 0.9000 | 4.6100 |
| O | W | 0.6690 | 4.2166 |
| COOH | COOH | 0.6500 | 3.0000 |
| COOH | W | 0.7627 | 4.5418 |

Full parameter files are available from:
https://github.com/CG-it/ffdb-sdk

Input files suitable for simulations with LAMMPS (Plimpton, 1995) with these parameter files can be prepared using the CG-it software:
https://github.com/CG-it/CG-it

**Hexatic Order Calculations.**

The average hexatic order of model bilayers was calculated using the final configurations from the corresponding simulations. The center position of each ceramide tail and each cholesterol molecule was estimated using the third aliphathic CG particle from its head group. Based on the lateral positions of each tail's six nearest neighbors, its local hexatic order ($|\psi_6|$) was then estimated by taking the norm of the bond orientation parameter ($\psi_6$) introduced by Halperin and Nelson (Halperin & Nelson, 1978). Cholesterol molecules were treated as nearest neighbors when calculating the $|\psi_6|$ values for the ceramide tails; however, the averages in Fig. S4 are based solely on the $|\psi_6|$ values of the ceramide tails and do not include $|\psi_6|$ values of the cholesterol molecules.

**Potential of mean force (PMF) and permeability calculations.**

Atomistic PMF calculations were initialized by inserting each small molecule at different depths on the pre-equilibrated 4×4 $nm^2$ hydrated bilayer. Lipid molecules with atoms within a 20 Å radius from the small molecule in the initial structure were translated along the bilayer in the away from the small molecule: the length of the translation was defined so as to remove all clashes for the nearest lipid neighbors, and linearly decreased to 0 for the lipids beyond 20 Å from the small molecule.

40 replicas for each system were simulated, each covering a 4 Å interval around the initial depth. Uniform coverage of each interval was achieved by using half-harmonic boundary restraints (also called a flat-bottom potential) outside the interval; the boundary forces were not included in the PMF. The centers of consecutive intervals were spaced by 1 Å, ensuring significant overlap between neighboring replicas. After minimization and equilibration in the $NP_xP_yP_zT$ (fully isotropic) ensemble for 0.3 µs, each PMF calculation was carried out for a total sampling time of 1 µs for each molecule (including all replicas). Sampling was enhanced by the Adaptive Biasing Force (ABF) method (Henin et al., 2010) as implemented in the Colvars module (Fiorin et al., 2013), by measuring free-energy gradients via projection onto the collective variable of the internal forces and integrating to produce the PMFs shown in **Fig. S8**.

Local maxima and local minima of each PMF were defined automatically by fitting quadratic curves to the PMF. The permeability of the outer bilayer leaflet was defined as:

$$k_P^{\text{[leaflet]}} = \left( \int \frac{\exp\left(\frac{G(z)}{k_\text{B}T}\right)}{D(z)} \mathrm{d}z \right)^{-1} \approx \exp\left(-\frac{G^*}{k_\text{B}T}\right) D$$

This approximation is justified by the presence of a single barrier with free-energy G* in each PMF, the exponential dependence of the expression on the free-energy, and the fact that the diffusion coefficient $D(z)$ is approximately constant along the lamella's normal.

#### Water exchange events and water droplet cluster analysis

The water exchange data in **Tables 1** and **2**, and related droplet sizes, were obtained by performing cluster analysis on the water particles in the configurations from the corresponding CG and AA simulations.  For the CG simulations, the cluster analysis was performed using a distance of 0.66 nm to extract nearest-neighbor water particles. Note that 0.66 nm corresponds to the first minimum in the radial distribution function for the CG water particles (Shinoda et al., 2007). In turn, CG water particles separated by less than 0.66 nm were labelled as part of the same cluster. For atomistic water, a distance cut-off of 0.35 nm was instead used to establish adjacent (i.e., connected) water molecules. Isolated water particles and clusters contains fewer than 10 CG water beads (i.e., 30 water molecules) were labelled as “free” water; this ensured that small transient clusters were not misidentified as droplets. Note that the smallest droplet size observed in the two lamellae CG simulation corresponded to 52 CG water beads (i.e., 156 water molecules). The water layers were easily identified by their much larger size compared to the water droplets.  In turn, water exchange events were detected by comparing a simulation’s successive configurations, and monitoring the movement of water particles between droplets, layers, and free states.

### SUPPLEMENTARY FIGURES

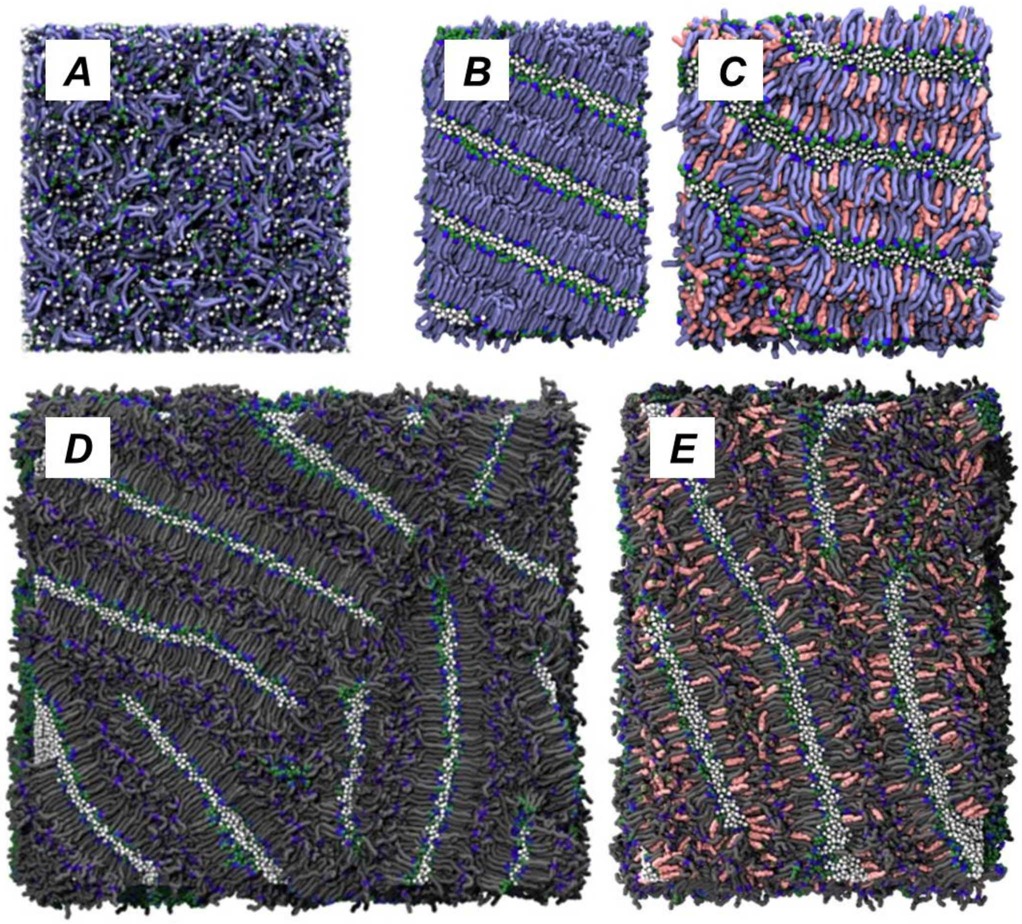


**Figure S1. Self-assembled lamellar structures predicted by MD simulations of ceramide mixtures with the CG model. (A)** Short-chain CER[NS] molecules (light blue) were initially prepared at randomized positions (white spheres are water particles) and allowed to self-assemble, with a lamellar structure as final result **(B)**. **(C)** Self-assembled lamellar structure of a 1:1 mixture of CER[NS] and cholesterol (pink). **(D,E)** Self-assembled structures of long-chain CER[EOS] molecules (gray) as a pure phase (D) and in a 1:1 mixture with cholesterol (E).

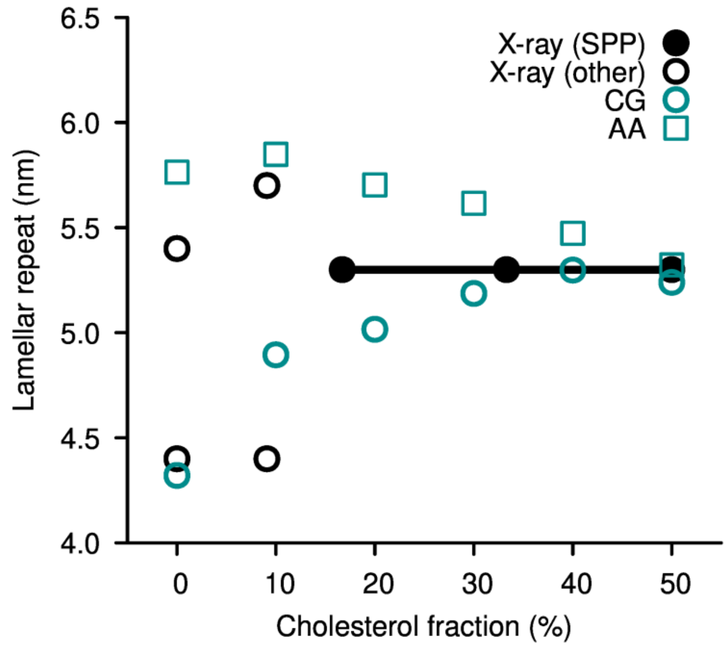


**Figure S2. Lamellar repeat lengths of CER[NS] model mixtures with increasing cholesterol content.** Black circles indicate values measured from X-ray diffraction (Mojumdar et al., 2015); depending on whether they were assigned to the SPP or other phase, these are shown as filled or empty, respectively. Blue symbols indicate values from simulations of small bilayers with the CG model (circles) and with the atomistic model (squares). Bilayer simulations were carried out in a water buffer (20:1 water:lipid ratio) for 0.2 μs. Lamellar repeat lengths were estimated by subtracting the excess water thickness (approximately 0.8 nm) from the mean unit cell spacing; the same protocol was used for data shown in Fig. S3.

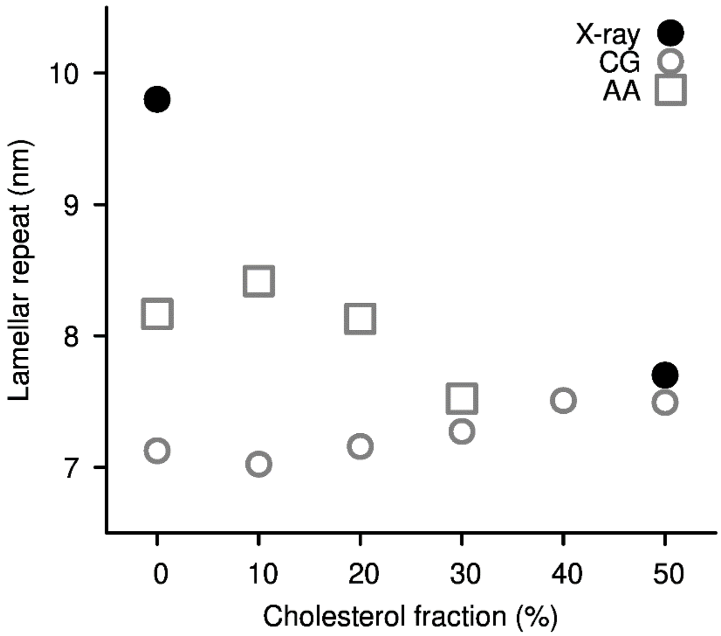


**Figure S3. Lamellar repeat lengths of CER[EOS] model mixtures with increasing cholesterol content.** Black circles indicate values measured from X-ray diffraction (Groen et al., 2010) and grey symbols values computed from simulations of small bilayers with the CG model (circles) and with the atomistic model (squares). The 9.8 nm repeat length of pure CER[EOS] (Groen et al., 2010) could not be reproduced with either simulation model.

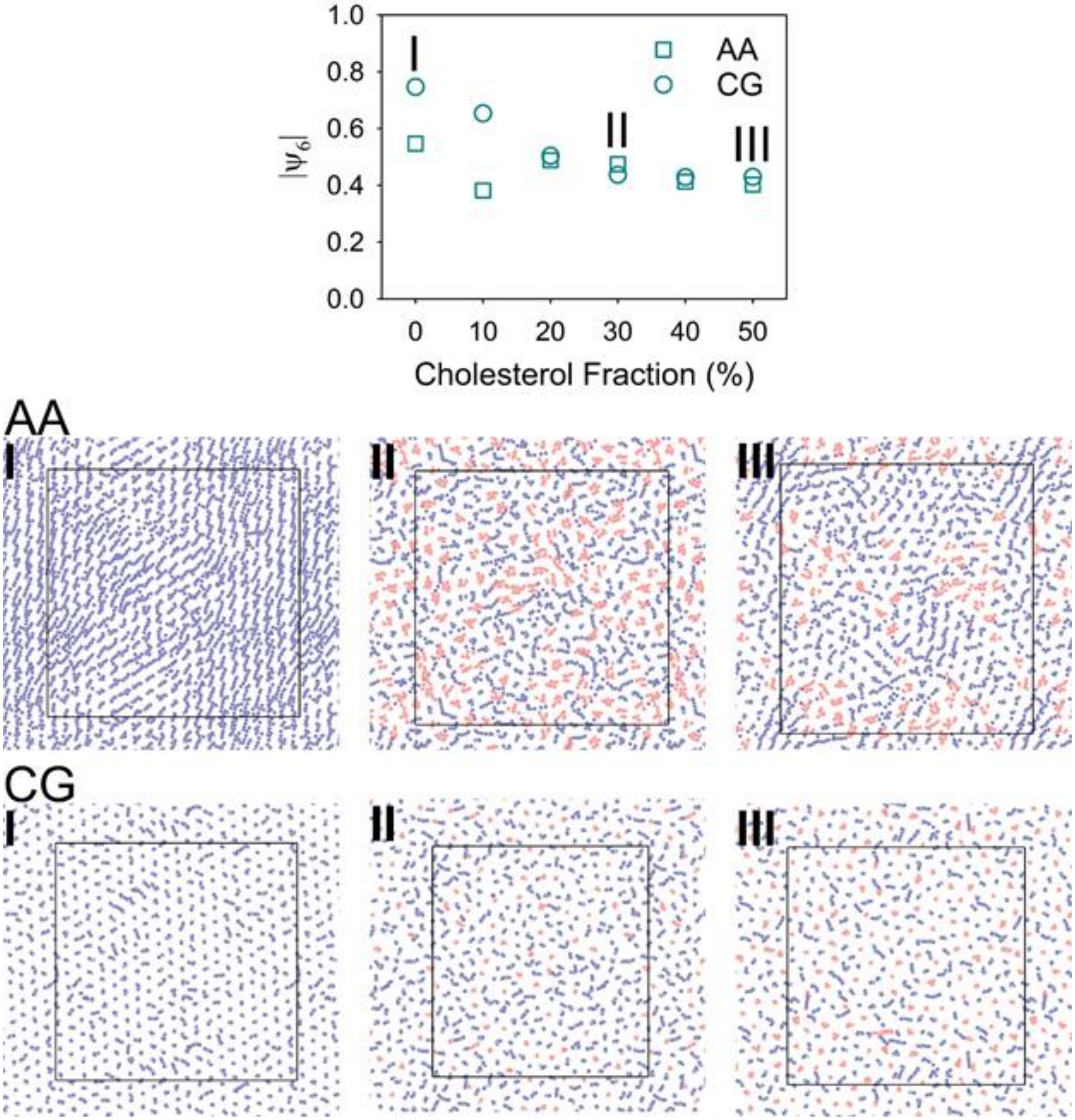


**Figure S4. Hexatic order of CER[NS] bilayers with increasing cholesterol fraction.** The top graph provides the variation in average local hexatic order for the ceramide tails in both the AA and CG simulations. The lower images provide snapshots of a leaflet from each system at cholesterol fractions of 0% (I), 30% (II), and 50% (III). Average positions over a 0.1 ns window are shown for visual clarity.

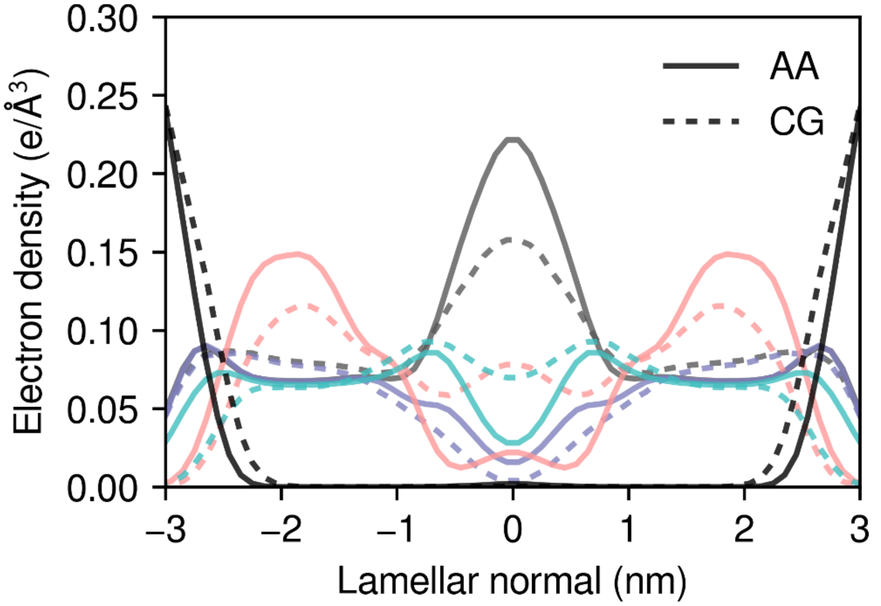


**Figure S5. Computed electron density profiles (EDPs) of skin lipid bilayers.** EDPs are plotted separately for CER[EOS], CER[NS], cholesterol and FFA; colors for each lipid type are as in other figures, while the water EDP is shown in black. Dashed lines indicate results from the CG simulations, and solid lines from a 1.5 μs atomistic simulation. For each lipid, the absolute value of the difference between atomistic and CG distributions, $|\rho_{AA} - \rho_{CG}|$, is less than 20% of the total density, indicating good agreement between atomistic and CG simulation models with independent initial conditions.

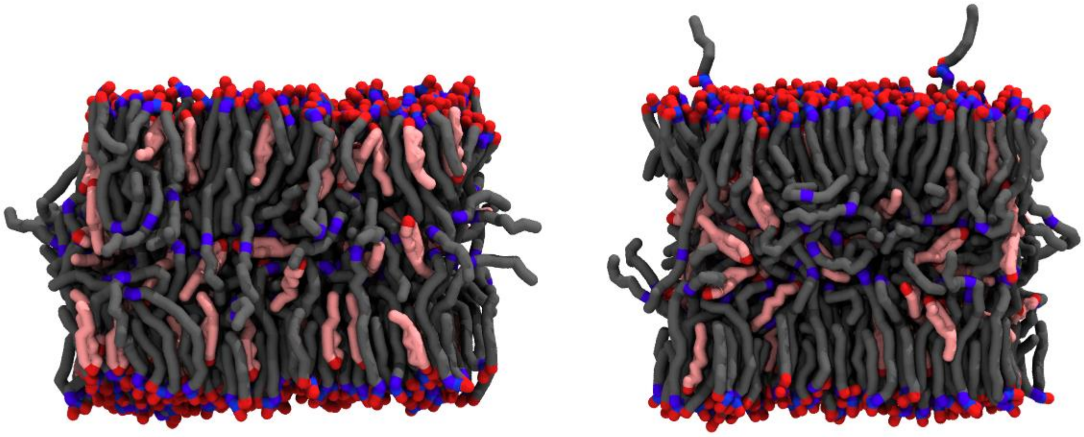

**Figure S6. Final bilayer snapshots of CER[EOS] mixtures with 50% cholesterol simulated with the CG model.** Water:lipid ratios are 20:1 in (left panel) and 5:1 (right panel): in the latter snapshot, few CER[EOS] molecules are in extended form, extending across the water layer. This feature is observed for both the 40% and 50% cholesterol mixtures, but not at lower cholesterol contents. Although the lamellar thicknesses are similar between 20:1 and 5:1 water:lipid, the latter appears to promote a tighter packing order of the saturated chains in the outer leaflet, and the movement of cholesterol to the region immediately under it, consistent with the position assumed in the LPP (Mojumdar et al., 2013).

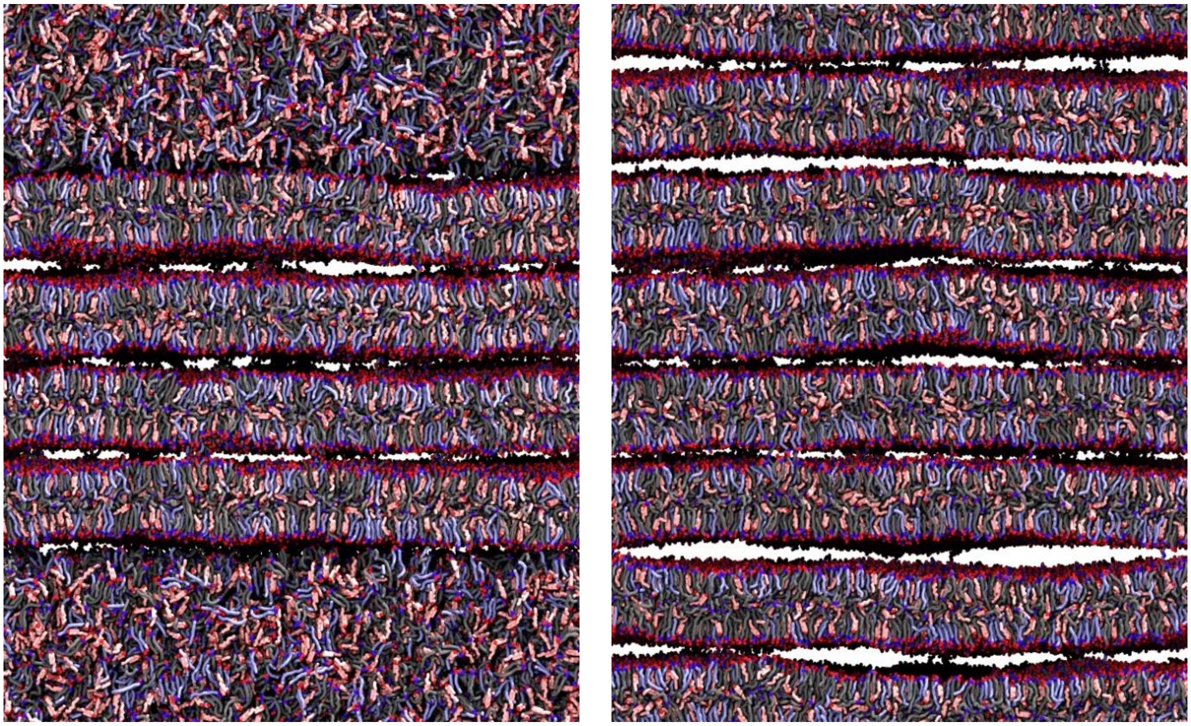

**Figure S7. Initial and final snapshots of a 2 μs-long simulation of a mixture of CER[EOS], CER[NS] and cholesterol in a 1:1:2 ratio at 5:1 water:lipid.** Lipid molecules are colored as in other figures, and water particles are hidden for clarity. 64,000 lipids were initially assembled in a lamellar phase (central four layers) and other 64,000 lipids in a randomized mixture. At the end of the simulation eight homogeneous bilayers are formed (unit cell 55×55×69 $nm^3$).

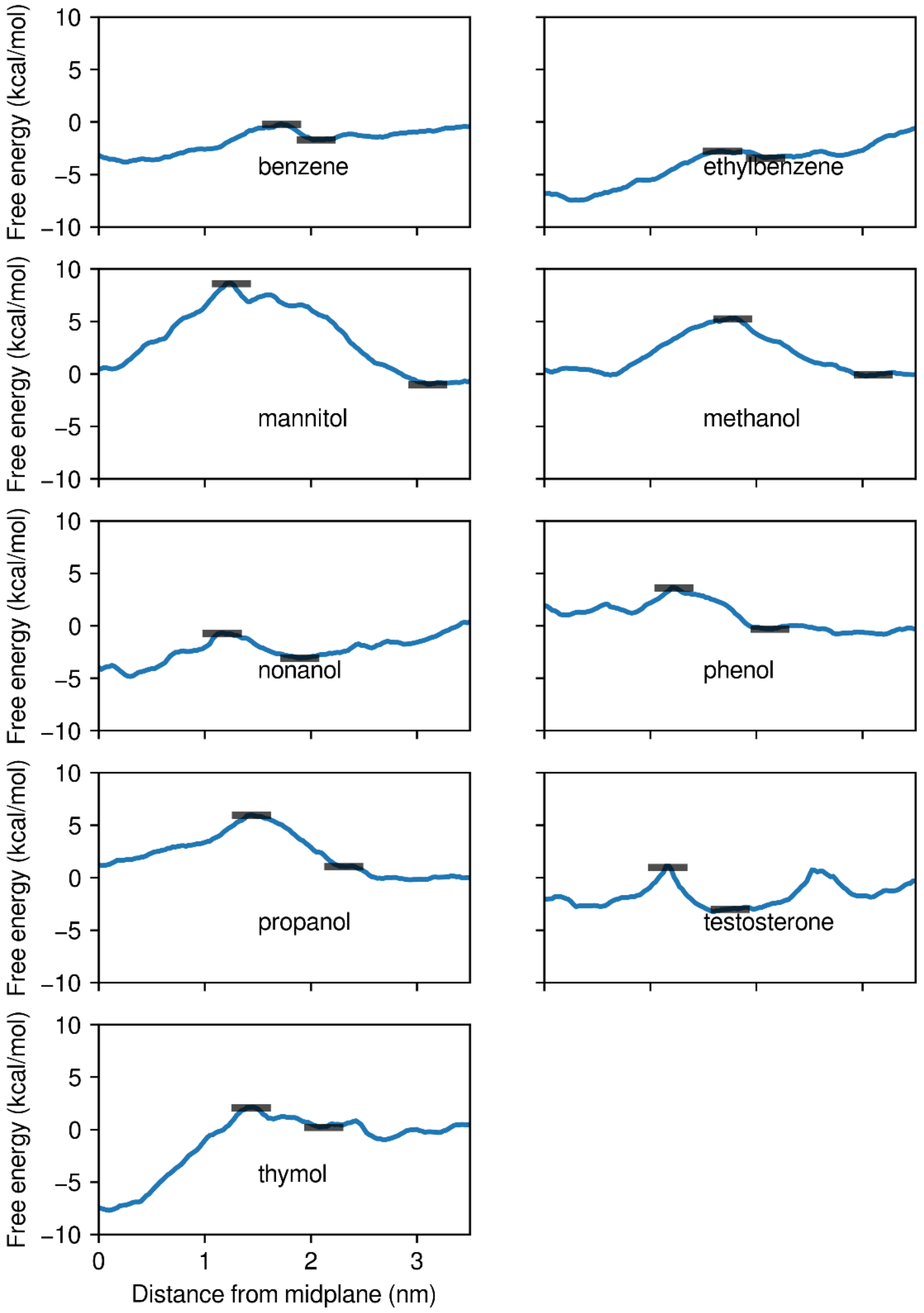


**Figure S8.** Shown are the potentials of mean force (PMFs) across a skin lipid bilayer rich in CER[EOS] using the adaptive biasing force (ABF) method with the thermodynamic-integration estimator applied to internal forces (Henin et al., 2010). For each PMF, black bands indicate the position of the highest free-energy point and the most favorable position in the model bilayer.

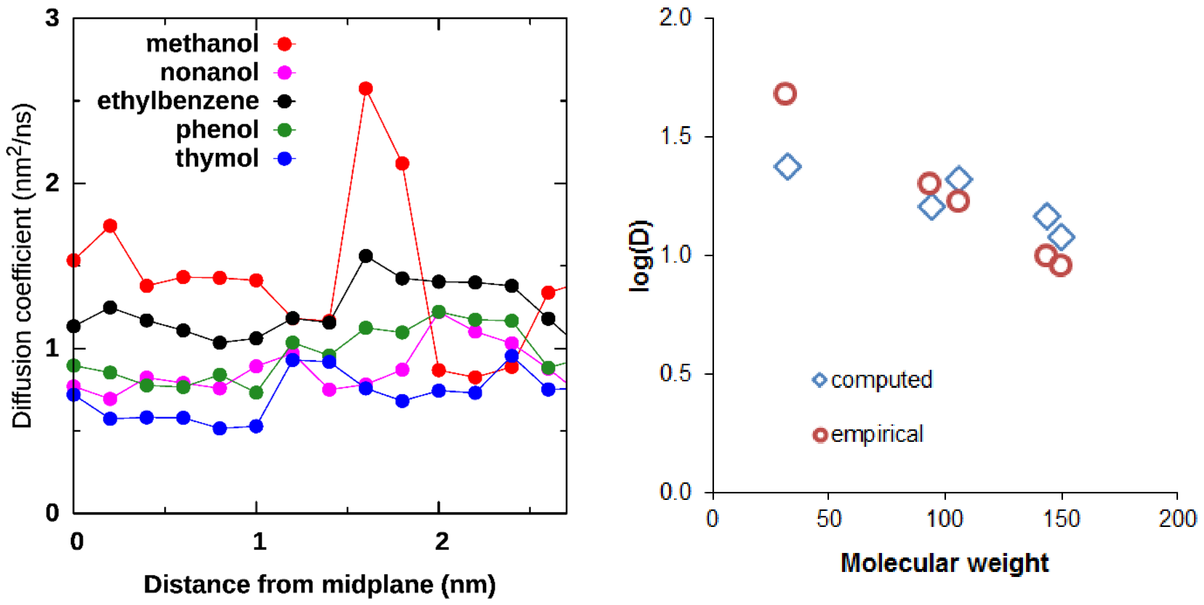


**Figure S9. (Left)** Shown are the diffusion coefficient profiles $D(z)$ across the SPP bilayer model on five molecules computed from simulation data using the procedure of Comer et al (Comer et al., 2013). **(Right)** Comparison between values of $D$ from simulation with those from the Potts-Guy equation (Potts & Guy, 1992); logarithms are in base 10, and the global multiplicative factor of the Potts-Guy model was scaled to allow comparison.

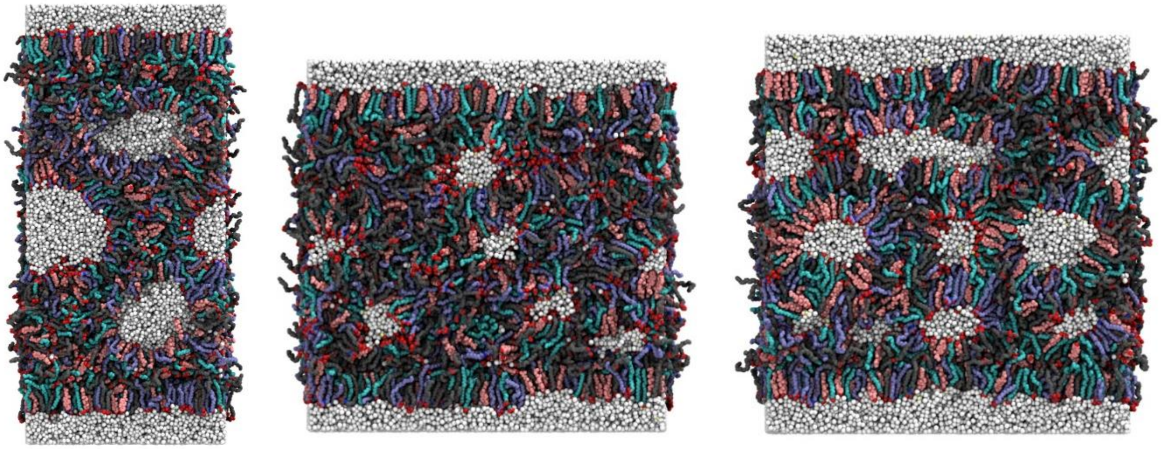

**Figure S10. (Left)** Final snapshot of the 16×16×32 $nm^3$ atomistic model at 5:1 water:lipid with fully-protonated FFAs. **(Center)** Final snapshot of the 24×24×32 $nm^3$ atomistic model at 2:1 water:lipid with mixture of protonated and unprotonated FFAs. **(Right)** Final snapshot of the 24×24×32 $nm^3$ atomistic model at 5:1 water:lipid with mixture of protonated and unprotonated FFAs.

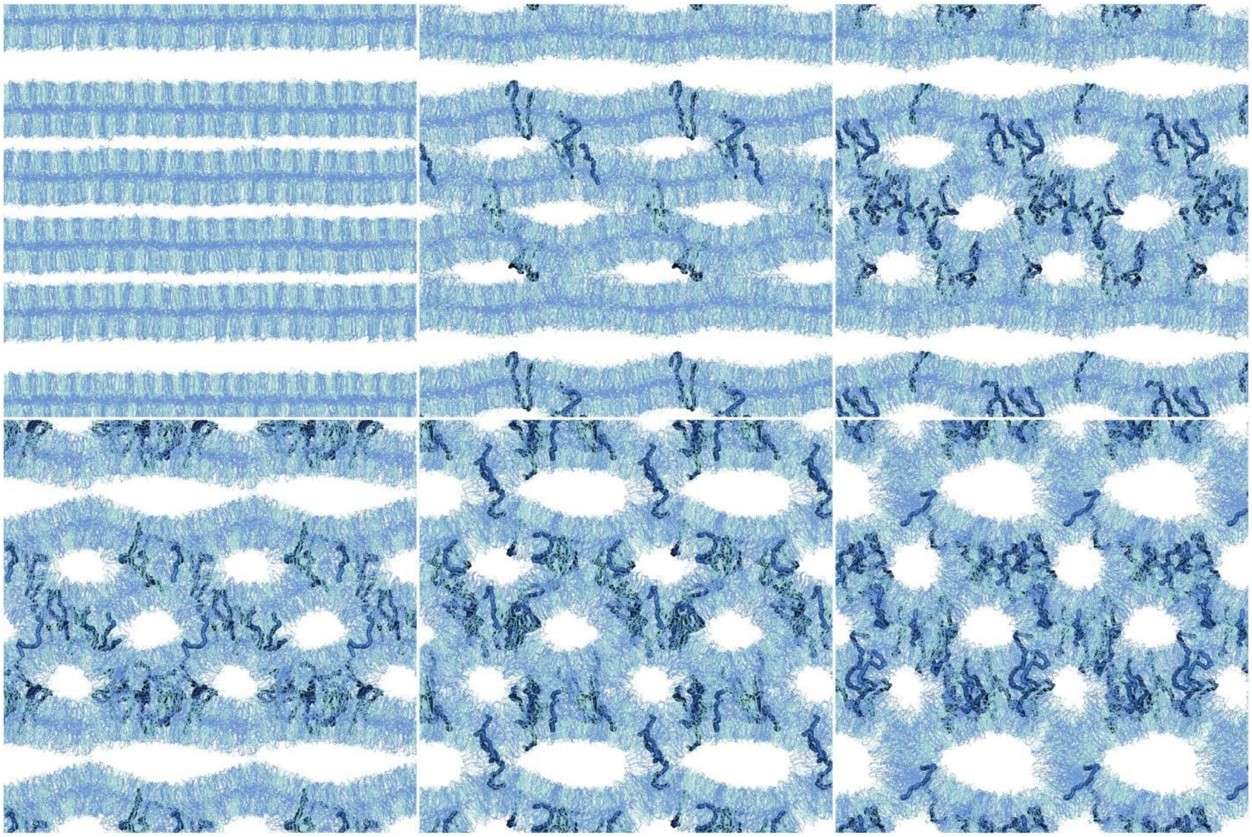

**Figure S11.** Shown are simulation snapshots every 50 ns during the heating stage (up to 250 ns) of the 16×16×32 $nm^3$ atomistic model with 5:1 water:lipid (**Fig. 2**). All lipids are shown as light blue; highlighted in darker blue are the CER[EOS] molecules with distance larger than 1.5 nm between the 18$^{th}$ carbon atoms of the sphingosine chain and fatty acid chain of the same molecule.

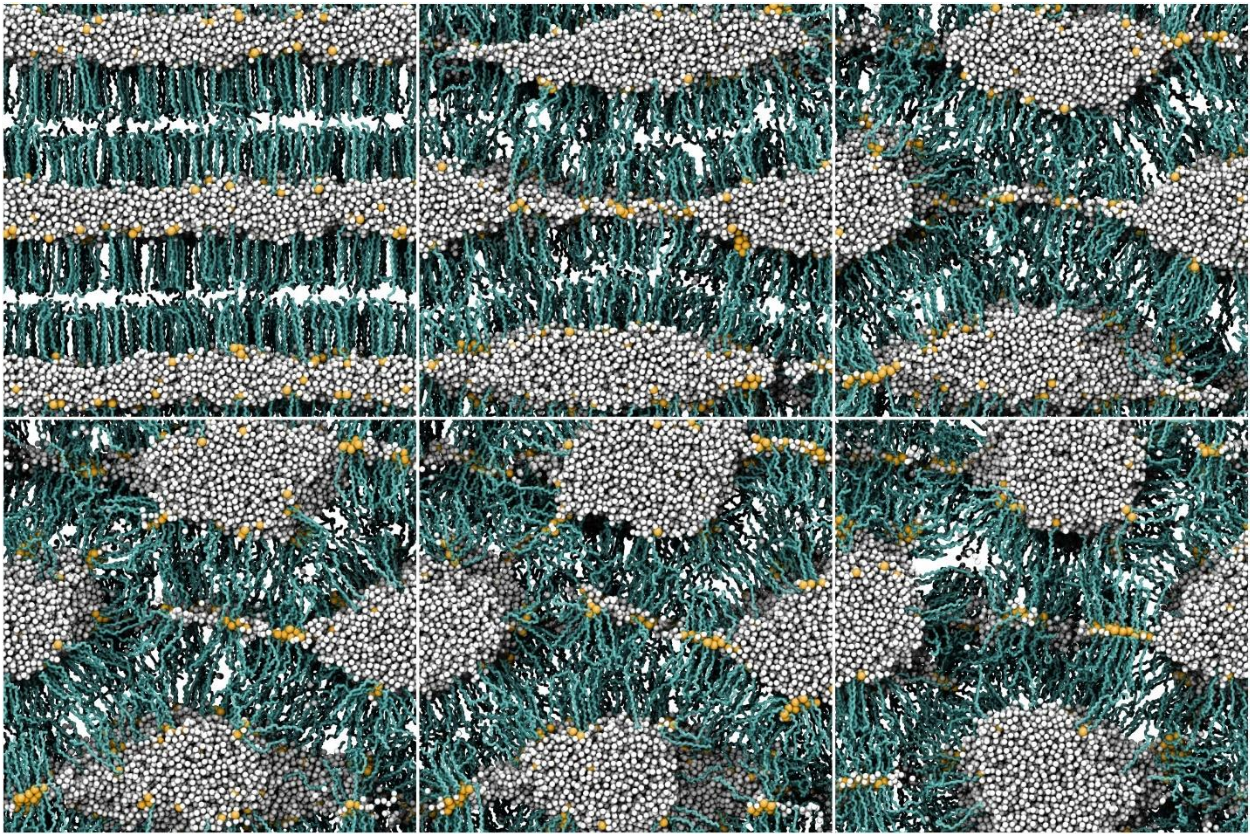

**Figure S12.** Snapshots of the same simulation shown in Fig. S11 highlighting the $Na^{+}$-mediated adhesion between unprotonated free fatty acids.

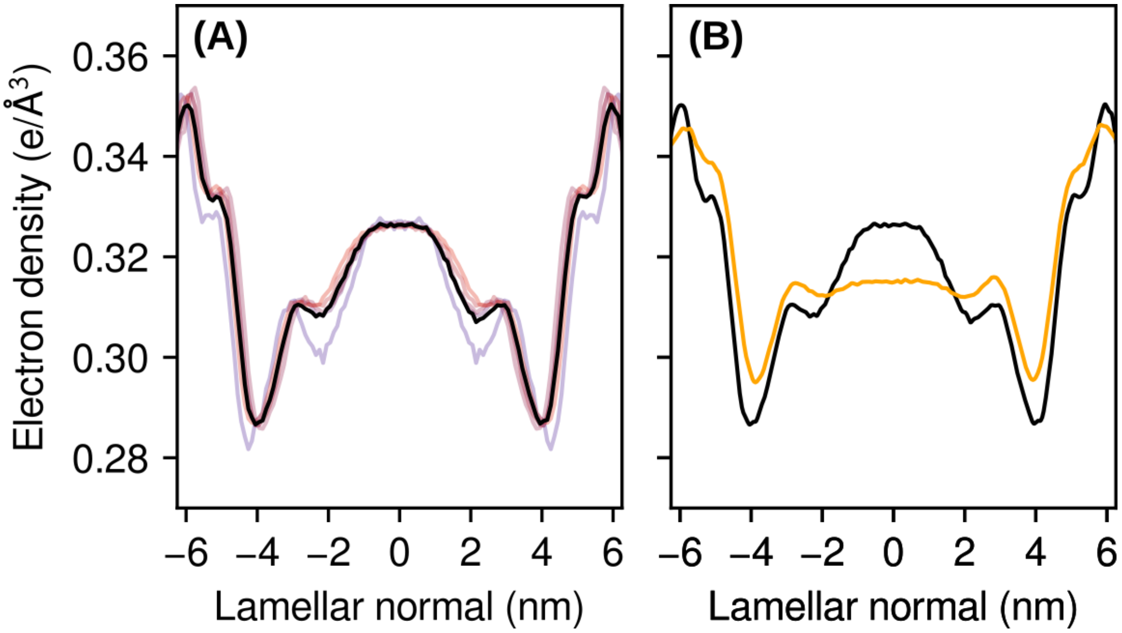


**Figure S13. (A)** Computed electron density profiles (EDPs) for the droplet-containing lamellae, shown as transparent lines (colors ranging from blue to red denote increasing droplet sizes), while the average EDP over all models is shown as a black line. Each EDP was computed over the last 5 μs of the corresponding CG simulation by assigning to the point position of each CG particle the corresponding number of electrons. **(B)** Comparison of the average EDP of droplet-containing lamellae in panel A (black) with the EDP from the dehydrated lamella (orange).

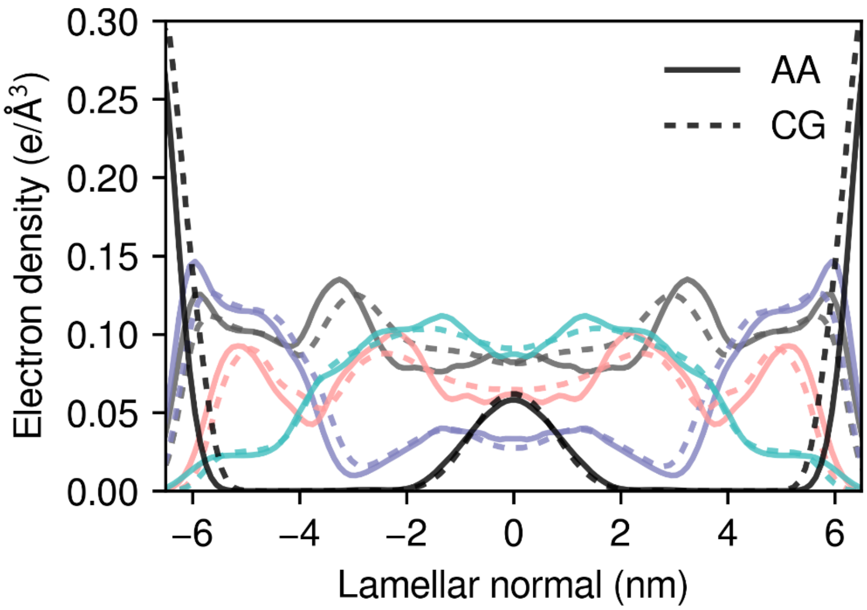


**Figure S14.** Computed electron density profiles (EDPs) are plotted separately for CER[EOS], CER[NS], cholesterol and FFA for the hydrated lamellae. Colors for each lipid type are as in other figures, while the water EDP is shown in black. Dashed lines indicate results from the CG simulations, and solid lines from atomistic simulations.

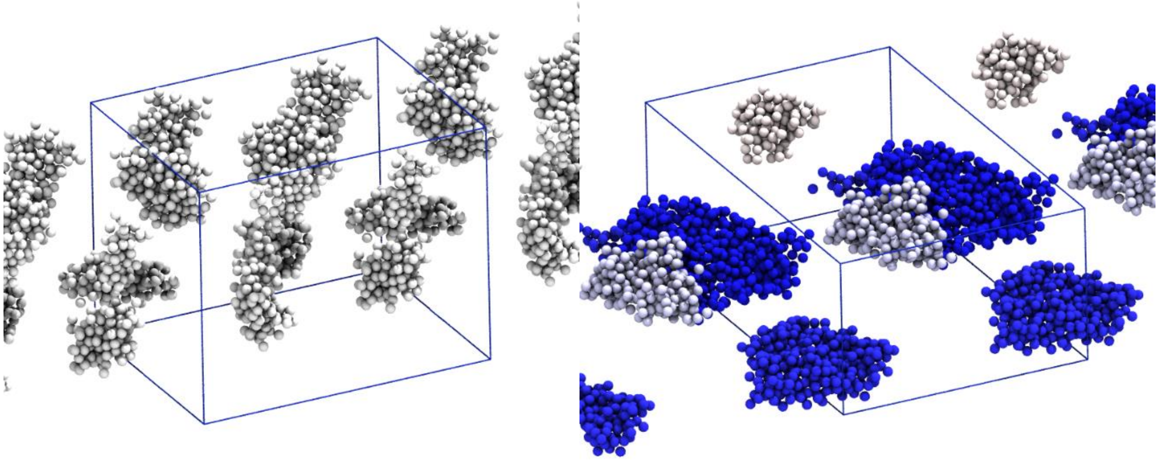

**Figure S15.** Shown are the initial (left) and final (right) snapshot of one simulation of a 3D inverse-micellar lipid unit cell: only water molecules are shown for clarity. In less than 10 μs, 6 of the initial 8 droplets coalesce into pseudo-lamellar domains (highlighted in blue), with 2 approximately spherical droplets remaining.

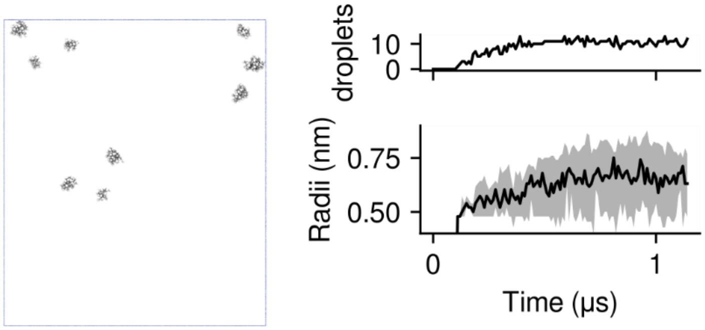


**Figure S16.** Simulation of the dehydrated lamella (**Figure 3B**) with permeation of water permitted; 9 droplets nucleate (left) but their growth is slowed down significantly (right) compared to the initial slab (**Figure 3A**) due to lower exchange of lipids between the interior and the outer leaflets.

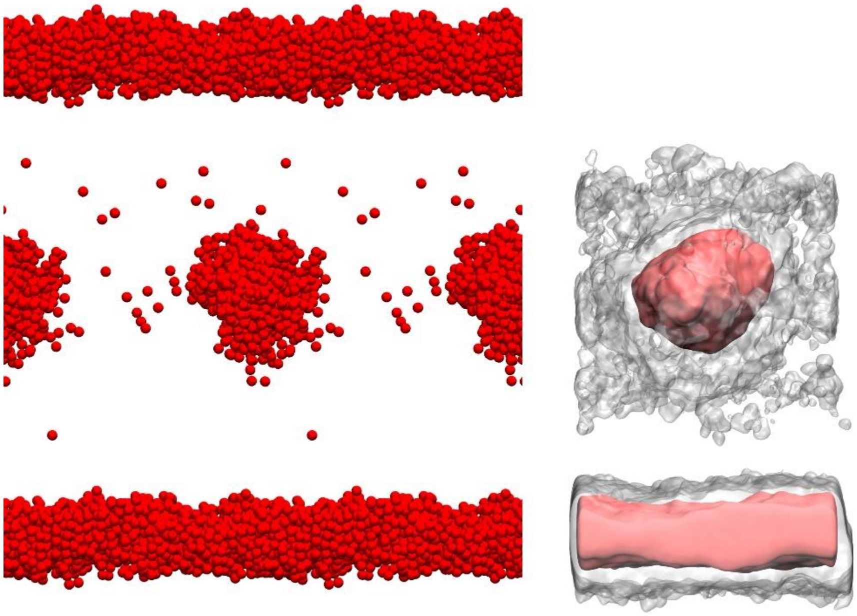

**Figure S17. Distribution of water molecules in the periodic atomistic simulation of an interstitial droplet.** On the left one of the simulation snapshots is shown: red spheres correspond to the oxygen atoms of the water molecules. The rest of the system is not shown for visual clarity.  On the right, the isosurfaces of water molecule oxygen number densities as averaged across the simulation. The resulting map had a number density ranging from 0 to ~0.27 atoms/Å$^3$. The red and the translucent surfaces correspond isovalues of 0.08 and 0.0008 atoms/Å$^3$, respectively.

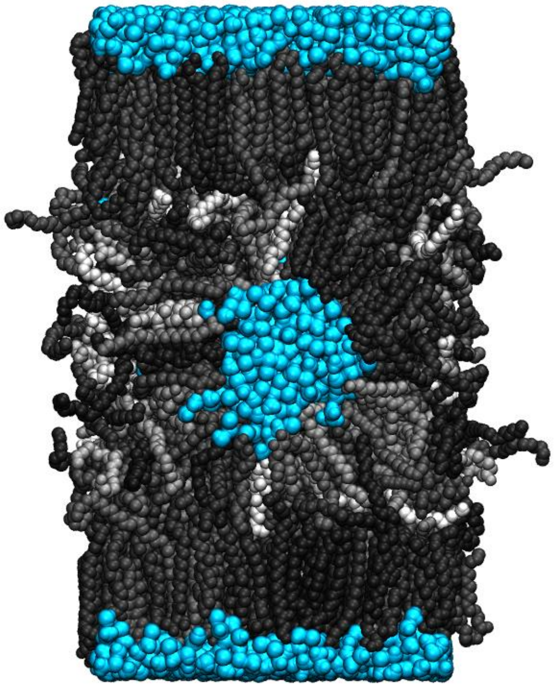

**Figure S18. Movement of lipids in the atomistic simulation of an interstitial droplet.** Shown is the final snapshot ($t$ = 2.2 μs) of the simulation, with water oxygens in light blue and each lipid molecule colored based on its displacement from the initial structure, ranging from 1 nm (black) to 8 nm (white); the latter also corresponds to the lateral dimensions of the unit cell. The first quartile, median and third quartile of the displacements' distribution are 1.3, 1.9 and 2.8 nm (in a CG simulation segment of equal length, these are 11, 20 and 33 nm, respectively).

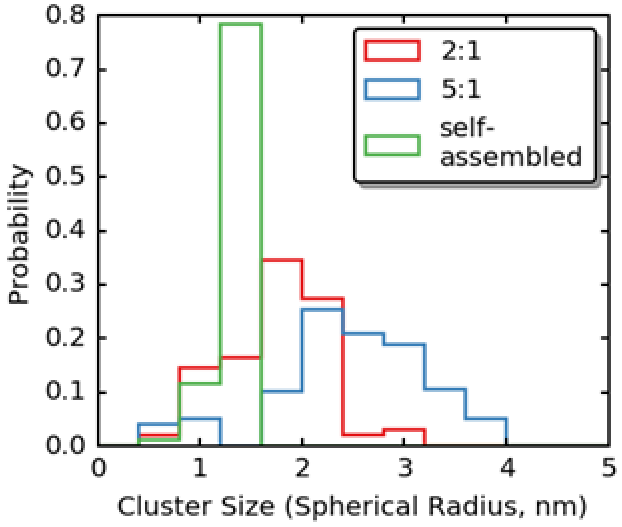


**Figure S19.** Shown is the distribution of droplet sizes (modeled as spherical for simplicity) resulting from the atomistic simulations in **Fig. 2** (red, blue) with the distribution of droplets self-assembled from individual water molecules in CG simulations of **Fig. 3** (green). Droplets with radii less than or equal to the optimal radius (1.3 nm) are approximately spherical, and their distributions follow similar profiles across the three data sets.

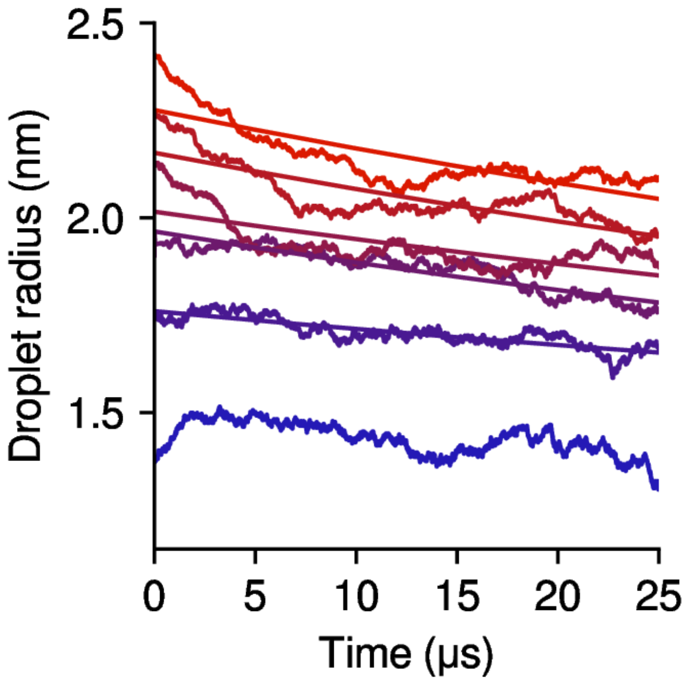


**Figure S20.** Time evolution of water droplets of different size. For the simulations that exhibit a significant drift, best-fit exponential lines are shown using the energy minimum predicted from Eq. 2 ($r^*$ = 1.3 nm) as a fixed parameter, and the initial radius and decay time as adjustable parameters. The resulting decay rates are plotted in **Fig. 4E**.

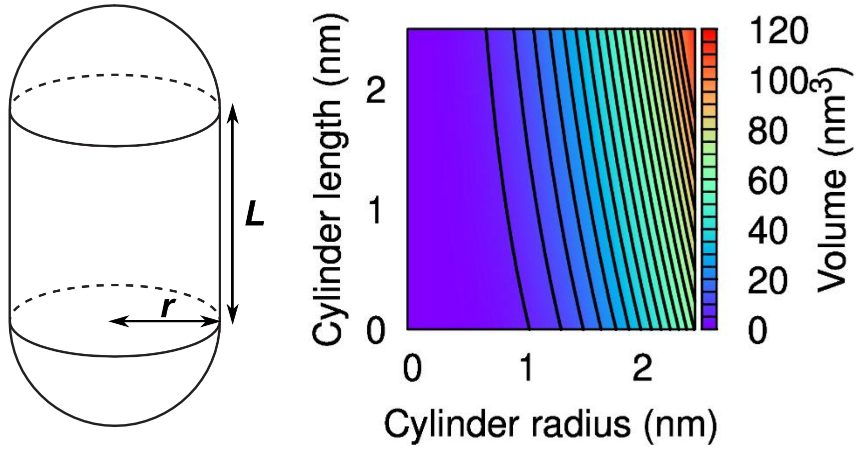


**Figure S21**. (**Left**) Schematic of the capsule used to model elongated intra-lamellar water droplets. (**Right**) Volume of the capsule as a function of the cylinder's radius *r* and length *L*.

### SUPPLEMENTARY REFERENCES


Comer, J., Chipot, C., & Gonzalez-Nilo, F. D. (2013). Calculating Position-Dependent Diffusivity in Biased Molecular Dynamics Simulations. *Journal of Chemical Theory and Computation, 9*(2), 876-882. doi:Doi 10.1021/Ct300867e

DeVane, R., Shinoda, W., Moore, P. B., & Klein, M. L. (2009). Transferable Coarse Grain Nonbonded Interaction Model for Amino Acids. *Journal of Chemical Theory and Computation, 5*(8), 2115-2124. doi:Doi 10.1021/Ct800441u

Fiorin, G., Klein, M. L., & Henin, J. (2013). Using collective variables to drive molecular dynamics simulations. *Molecular Physics, 111*(22-23), 3345-3362. doi:10.1080/00268976.2013.813594

Groen, D., Gooris, G. S., & Bouwstra, J. A. (2010). Model Membranes Prepared with Ceramide EOS, Cholesterol and Free Fatty Acids Form a Unique Lamellar Phase. *Langmuir, 26*(6), 4168-4175. doi:10.1021/la9047038

Halperin, B. I., & Nelson, D. R. (1978). Theory of Two-Dimensional Melting. *Physical Review Letters, 41*(2), 121-124. doi:10.1103/PhysRevLett.41.121

Henin, J., Fiorin, G., Chipot, C., & Klein, M. L. (2010). Exploring Multidimensional Free Energy Landscapes Using Time-Dependent Biases on Collective Variables. *Journal of Chemical Theory and Computation, 6*(1), 35-47. doi:Doi 10.1021/Ct9004432

Mogurampelly, S., MacDermaid, C. M., Percec, S., Klein, M. L., & Fiorin, G. (2019). Aggregation of poly($p$-phenylene terephthalamide) chains: Emergence of fiber defects. *Physical Review Materials, 3*(1), 015602. doi:10.1103/PhysRevMaterials.3.015602

Mojumdar, E. H., Gooris, G. S., & Bouwstra, J. A. (2015). Phase behavior of skin lipid mixtures: the effect of cholesterol on lipid organization. *Soft Matter, 11*(21), 4326-4336. doi:10.1039/c4sm02786h

Mojumdar, E. H., Groen, D., Gooris, G. S., Barlow, D. J., Lawrence, M. J., Deme, B., & Bouwstra, J. A. (2013). Localization of cholesterol and fatty acid in a model lipid membrane: a neutron diffraction approach. *Biophysical Journal, 105*(4), 911-918. doi:10.1016/j.bpj.2013.07.003

Plimpton, S. (1995). Fast Parallel Algorithms for Short-Range Molecular-Dynamics. *Journal of Computational Physics, 117*(1), 1-19. doi:DOI 10.1006/jcph.1995.1039

Potts, R. O., & Guy, R. H. (1992). Predicting Skin Permeability. *Pharmaceutical research, 9*, 663-669.

Shinoda, W., Devane, R., & Klein, M. L. (2007). Multi-property fitting and parameterization of a coarse grained model for aqueous surfactants. *Molecular Simulation, 33*(1-2), 27-36. doi:Doi 10.1080/08927020601054050
